# The phenotypic landscape of the model firmicute *Bacillus subtilis*

**DOI:** 10.64898/2026.05.13.724699

**Authors:** Lili M. Kim, Byoung-Mo Koo, Hannah N. Burkhart, Daniela Vollmer, Waldemar Vollmer, Horia Todor, Carol A. Gross

## Abstract

Firmicutes are gram-positive bacteria with important roles in human health, disease, and industry. However, more than a quarter of genes in the model firmicute *Bacillus subtilis* remain completely uncharacterized, including numerous core phylum-specific genes. Here, we design a compact pooled CRISPRi library targeting all protein-coding genes in *B. subtilis* and test its growth in ∼150 stress conditions. Using data from this screen as a hypothesis generator, we perform targeted experiments, revealing that the conserved essential firmicute protein YneF is part of the SRP co-translational protein secretion pathway. We also demonstrate that ECF-transporters play a previously unknown but broadly conserved role in cell wall homeostasis, perform an unbiased analysis of amino acid crossfeeding, and make additional discoveries about bacterial competition. In addition to these major contributions to our understanding of *B. subtilis* biology (and gram-positive firmicutes in general), this work provides a rich dataset that will nucleate future studies of uncharacterized genes and presents a framework for accessible full-genome functional genomic screens in other bacteria.

**SIGNIFICANCE:** Large-scale chemical genomics screens facilitate the characterization of genes by generating phenotypic data across a library of gene mutants. Here, using CRISPRi, we designed a compact pooled library targeting all genes in *B. subtilis* and screened growth of the library in ∼150 conditions. Importantly, we used this dataset to expand *B. subtilis* biology on levels ranging from molecular pathways to bacterial communities. We discovered a role for an essential firmicute gene in protein secretion, implicated two new players in cell wall homeostasis, and unraveled fundamental factors driving competition and cooperation in the soil. This rich dataset will serve as a hypothesis generator, expanding our set of bacterial phenotypic data and driving future experiments in *B. subtilis* and other gram-positive firmicutes.

## INTRODUCTION

*Bacillus subtilis* is an important microbe for biomedical research and industrial applications. It is the model organism for gram-positive firmicutes, which include numerous pathogens (e.g., *Staphylococcus aureus, Streptococcus pneumoniae, Clostridium difficile*) and commensals (e.g., *Holdemanella biformis, Ruminococcus bromii, Lactobacillus plantarum*), boasts powerful tools for genetic and molecular research (1–6), and is widely recognized as a workhorse in biotechnology (7, 8). However, despite being one of the most intensively studied organisms, the function of ∼25% of all *B. subtilis* genes remain uncharacterized or poorly understood (9). Such a gap can be addressed by large-scale chemical genomics screens, wherein the fitness of each strain in a genome-wide mutant library is measured in a wide range of stress conditions. These chemical genomic screens identify genes required in specific conditions, and, importantly, associate genes of unknown function to known pathways on the basis of correlated phenotypes across conditions (phenotypic signatures; (10)). Chemical-genomics screens have been previously performed in other firmicutes; however, these have been technically limited. Large genome-wide screens in *S. aureus* (11) and *S. pneumoniae* (12) used TnSeq (13, 14), which cannot query essential gene phenotypes and is limited by sequencing depth. Conversely, small scale CRISPRi screens in *B. subtilis* and *S. pneumoniae* targeted only essential genes and used an arrayed approach (4, 15). Moreover, all of these screens used a small selection of (primarily) antibiotic stressors (20-32 conditions), which limits the interpretability of correlated phenotypic signatures.

Here, we overcome these limitations and perform a comprehensive chemical genomics screen in *B. subtilis* using a pooled CRISPRi library targeting all protein-coding genes. Our library combines full knockdown of non-essential genes (approximately equivalent to their deletion) with graded partial knockdown of essential genes using mismatch-CRISPRi to titrate sgRNA efficacy (16), which allows probing their mutant phenotypes without cell death. By designing our library to maximize phenotypic data while minimizing library size, we facilitate screening and enable extensive multiplexing, reducing experimental effort and sequencing cost. We subjected our library to an extensive array of conditions (n = 149), which included antibiotics targeting all essential processes as well as environmental stressors such as pH changes, osmotic stress, and non-antibiotic chemical perturbations. Our final dataset yielded many novel phenotypes and gene-gene correlations, which: 1) revealed the role of the previously uncharacterized essential gene *yneF*; 2) discovered narrowly and broadly conserved players in cell wall homeostasis; and 3) provided systems-level overviews of inter- and intra-species interactions and crossfeeding. We present this robust dataset as a powerful new resource for understanding firmicute biology and as a framework for accessible full-genome functional genomic screens in other organisms.

## RESULTS AND DISCUSSION

### A genome-wide pooled CRISPRi screen in *B. subtilis*

Our pooled CRISPRi library targeted all protein-coding genes in *B. subtilis* using a previously described and well-established inducible CRISPRi system (4, 5, 16). Non-essential genes (n = 3,981 genes) were fully knocked-down by two fully complementary, computationally optimized sgRNAs (Methods; (17)), while essential genes and those with strong growth defects (n = 307 genes; (5)) were targeted by 12 sgRNAs spanning the full range of possible knockdown levels explored in a previous study ((16); Fig. 1A; Data S1; Methods). This compact library (n = 11,559 sgRNAs) is easy to handle and multiplex, facilitating screening in a large number of conditions.

**Fig. 1.**
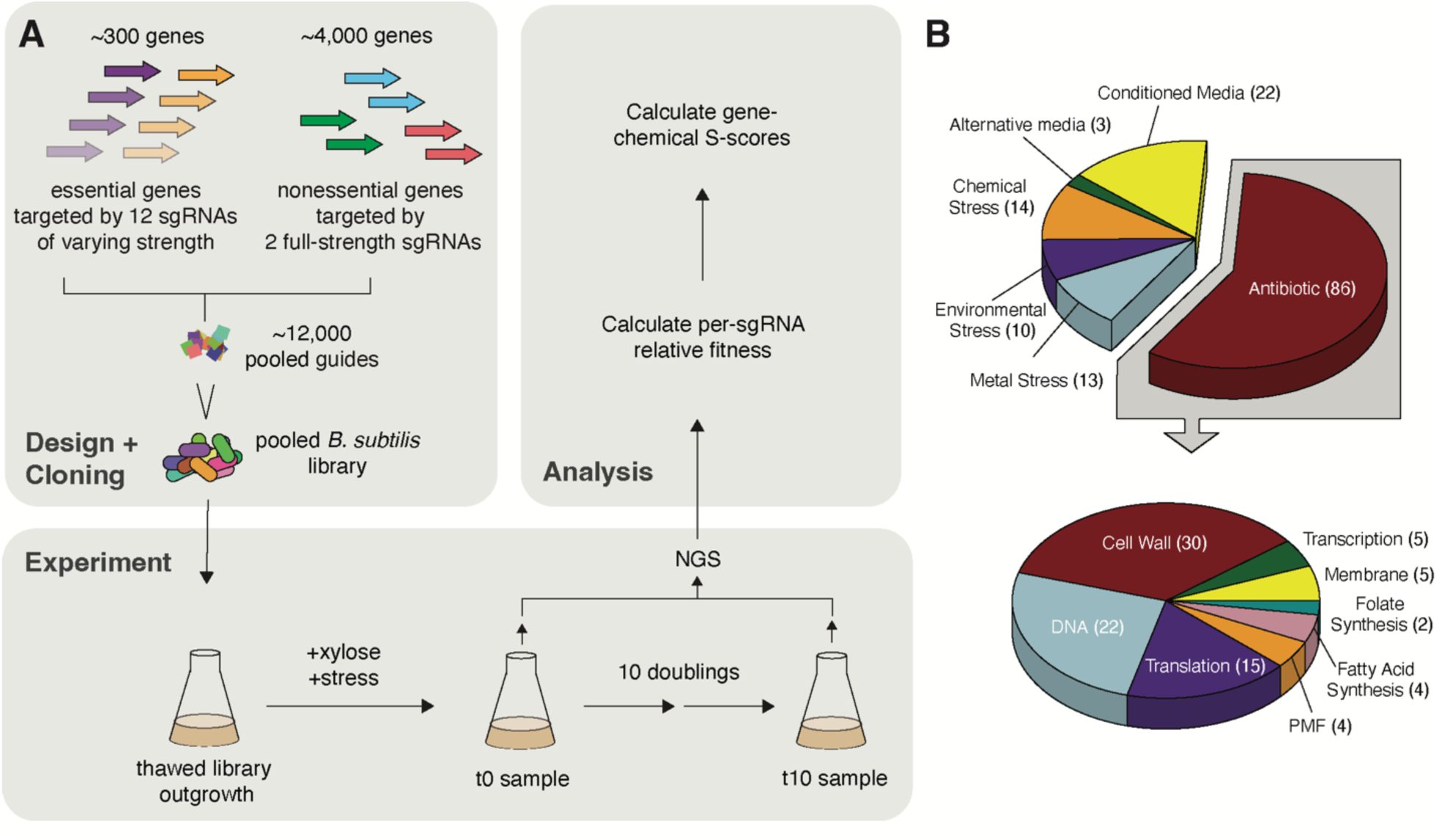
Design of full-genome CRISPRi chemical genomics screen. A. Strategy for design (top left), experimental setup (bottom), and analysis (top right) of an 11,559-sgRNA CRISPRi library targeting the full *B. subtilis* genome. Full details provided in Methods. B. Categories of stress conditions tested (top) and sub-categories of antibiotics tested (bottom). When possible, each stressor was tested at two sub-MIC concentrations (∼10% and 25% growth inhibition); each concentration of each stressor is counted.

We subjected our CRISPRi knockdown library to 83 unique stressors targeting all major processes in *B. subtilis,* including antibiotics (e.g., β-lactams, macrolides, fluoroquinolones), antimicrobials (e.g., ethidium bromide, NaCl, SDS), metals (e.g., ZnCl_2_, MnCl_2_, CuCl_2_), diverse environments, and others (Fig. 1B). Where possible, we used two sub-MIC concentrations of each stressor (∼10% and ∼25% growth inhibition) to maximize our ability to accurately measure both resistance and susceptibility (149 conditions). dCas9 expression was induced simultaneously with the addition of each stressor in exponentially growing cells. Cultures were sampled for next generation sequencing immediately before dCas9 induction and after 10 doublings post induction, as previously described (16, 18–20). We calculated the relative fitness (RF) of each strain in each condition by comparing its relative abundance before and after the experiment (Fig. 1A; Table S1). RF values were consistent across replicates (*r* > 0.95; Fig. S1A-B), and with previously published CRISPRi and knockout studies (*r* > 0.95; (5, 16), Fig. S1C-D, Methods).

We converted our RF values to chemical-gene interaction scores (S-scores; (21)), which facilitate the identification of strains with condition-specific growth differences and accounts for RF measurement variability (Methods). sgRNAs for the 307 essential/semi-essential genes were binned into low, medium, and high groups, with separate S-scores for each bin. Our final dataset consists of a matrix of S-scores for each of the 4,776 genes/bins by the 149 conditions that passed our quality-control criteria (Table S2; Methods), and contains 13,104 significant (|S-score| > 5) and 5,783 strong (|S-score| > 7) chemical-gene interactions, as well as many weaker interactions (Fig. 2, Table S2). Many of these recapitulated previously reported chemical-gene interactions: knockdown of *fabF* sensitized cells to cerulenin (4, 22), knockdown of *vmlR* sensitized cells to lincomycin (23), and the essentiality of *fpa* (*ylaN*) was rescued by excess iron (4, 24) (Table S2).

**Fig. 2.**
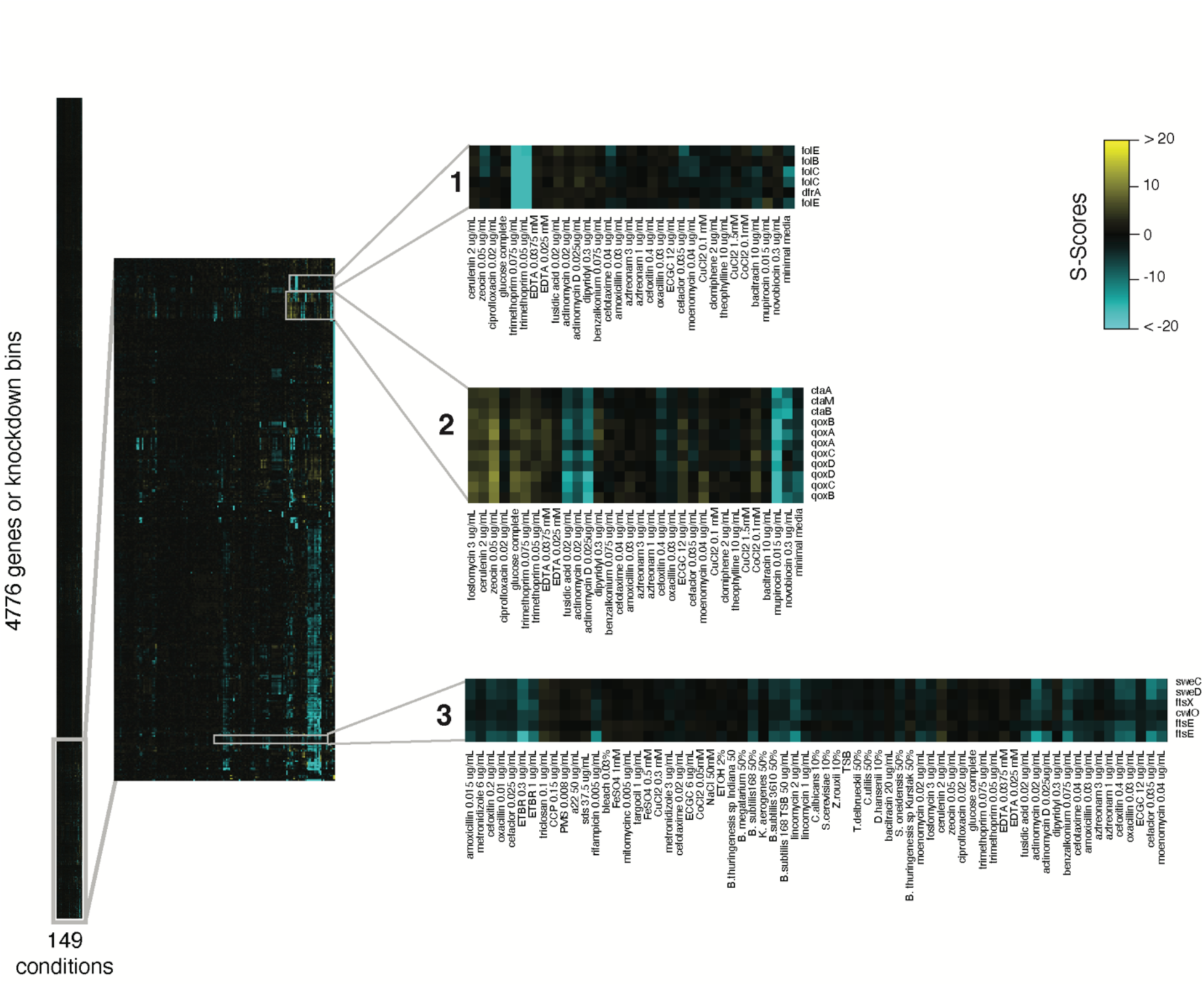
Clustered heatmap of fitness scores for 4776 nonessential genes or binned knockdowns of essential- or semi-essential genes in response to all 149 conditions. Zoomed insets demonstrate co-clustering of conditions (x-axis) and genes (y-axis): 1. *dfrA* and *folBCE*, essential biosynthesis genes of folate, co-cluster due to shared sensitivity to trimethoprim, which targets DfrA. 2. *qoxABCD,* the cytochrome aa3 oxidase, *ctaAB,* synthesis genes of the heme cofactor used by the Qox oxidase, and *ctaM,* required for Qox assembly and function(25), cluster together due to diverse positive (e.g. zeocin) and negative (e.g. mupirocin) phenotypes. 3. *cwlO,* involved in cell elongation, *ftsEX,* required for CwlO function, and *sweCD*, cofactors of FtsEX (26), cluster together due to shared sensitivity to cell-wall drugs. Essential genes appear multiple times in the heatmap because they have three separate sets of S-scores for low, medium, and high knockdown efficiency bins. The three bins may not always cluster together; high-knockdown bins are less likely to show specific negative phenotypes (as cells are already sick due to the strong knockdown of an essential gene), while low-knockdown bins are less likely to show specific positive phenotypes (as these mutants often grow similarly to WT).

Hierarchical clustering of our S-scores revealed genes with the most similar phenotypes (Fig. 2), and identified functional modules that span multiple operons. For example, the essential folate synthesis genes *dfrA* and *folBCE* cluster together, as expected (Fig. 2 inset 1). The terminal caa3 oxidase encoded by *qoxABCD*, which uses a heme A cofactor, clusters with the heme A biosynthesis genes (*ctaAB*) and *ctaM*, which is required for Qox activity ((25); Fig. 2 inset 2); demonstrating that related genes can have similar phenotypes beyond specific drug-target interactions. Finally, the peptidoglycan (PG) hydrolase *cwlO* clusters with its regulator, *ftsEX*, and with FtsEX cofactors *sweCD* ((26); Fig. 2 inset 3).

More broadly, phenotypic correlations were broadly indicative of functional connections: a majority of gene pairs with highly correlated phenotypes (*r* > 0.90, 674/1056; Table S3) had functional interactions documented in the STRING database (even when accounting for the polarity of CRISPRi knockdown by excluding intra-operon interactions; Fig. S3; Methods). Taken together, these analyses suggest that our data accurately reflects known gene-chemical and gene-gene interactions. Importantly, many interactions from our screen reflect completely new information: we found at least one strong phenotype for 18% of genes with function “unknown” in SubtiWiki ((27); Table S2).

### High-throughput chemical genomics expands previous studies

The genome-wide and unbiased nature of our data synergizes with previous genome-wide and targeted studies, allowing us to confirm and substantially expand results from previous studies in *B. subtilis* and other species, as illustrated by the three stories below.

We recently quantified genetic interactions (GIs) between cell-envelope genes in *B. subtilis* (28). These data can be analyzed together with chemical genomics data to better understand the mechanisms underlying observed GIs. For example, our GI screen implicated four genes (*yrrS* [*ragB* (29)]*, ytxG* [*facZ* (*30*)], *ypbE,* and *yerH*) in cell division due to their strong negative GIs with *ezrA,* a negative regulator of FtsZ-ring formation. Here, we find that knockdown strains of *yrrS/ragB, ytxG/facZ*, and *ypbE* were highly sensitized to moenomycin (S-scores < -24; Table S2), which inhibits the transglycosylation step of PG synthesis, while *yerH* knockdown strains were resistant to moenomycin (S-score = 10; Table S2). This suggests that despite shared GIs with *ezrA, yerH* may play a divergent role from *yrrS/ragB, ytxG/facZ,* and *ypbE*. Intriguingly, other cell division gene knockdown strains also were sensitized to moenomycin, including *ftsZ*, *minJ,* and *gpsB,* raising the possibility that other undercharacterized genes strongly sensitized to moenomycin (e.g., *yabM, ydbJKL,* or *yubA*) may also function in cell division.

Inhibition of fatty acid synthesis (FAS) was previously shown to rescue growth of *B. subtilis* lacking the central PG synthase *ponA* (31). Our data significantly expands this finding, demonstrating that knockdown of other elongasome genes, including essential ones like *rodA* and *mreC,* is also rescued by limiting FAS (Fig. S2A). Moreover and conversely, we found that inhibiting FAS was lethal in mutants with impaired divisomes, reinforcing the important connection between membrane synthesis and cell wall surface area and raising new questions about co-regulation of these processes (Fig. S2A).

Finally, our data extends recent findings in gram-negative bacteria that link the stringent response to Fe-S cluster synthesis to gram-positive bacteria. Damage to Fe-S clusters, required for synthesis of branched- and sulfur-containing amino acids, triggers the stringent response in *Salmonella enterica, Enterobacter cloaceae,* and *Klebsiella pneumoniae* (32). We found that knockdown of the *suf* operon (involved in Fe-S cluster synthesis) sensitizes *B. subtilis* to mupirocin (Fig. S2B), which stops cell growth by inhibiting IleS and triggering the stringent response. Moreover, we found that depleting *mntR* (which normally represses Mn^2+^ import (33)) also sensitizes *B. subtilis* to mupirocin, likely because high Mn^2+^ disrupts the function of Fe-S cluster-containing proteins ((32); Fig. S2B). Our results orthogonally confirm Fe-S damage as an input to the stringent response, which is exacerbated by mupirocin-induced amino acid limitation, and extend this connection to gram-positive bacteria.

These examples demonstrate how integrating our data with targeted and high-thoughput experiments from *B. subtilis* and other bacteria allows it to transcend *B. subtilis* biology, verifying and expanding regulatory themes, generating hypotheses, and revealing cross-organism similarities.

### YneF is a conserved, essential component of the SRP

*yneF* is an essential and conserved gene across Firmicutes (5) and Tenericutes (34–36) and has been flagged as an understudied, highly expressed gene in *B. subtilis* (9). Structural predictions by AlphaFold3 suggest that YneF contains a transmembrane helix and may dimerize (assessed by interface predicted template modeling score [ipTM], the accuracy score of a predicted complex; ipTM = 0.47, Fig. S4A). Our data revealed strong phenotypic correlations between *yneF,* and genes encoding proteins involved in the Signal Recognition Particle (SRP) pathway which performs co-translational protein insertion (Fig. 3A-B). The SRP machinery, universally conserved in all domains of life, minimally consists of 4.5S/6S/7SL RNA (in *B. subtilis*, a 6S RNA encoded by *scr*), the SRP54/Ffh protein, and the signal receptor (SR, FtsY). The SRP (6S and Ffh) binds signal sequences as they emerge from the ribosome and delivers the ribosome to SR (FtsY), triggering GTP hydrolysis and conformational changes that enable binding to the Sec (SecYEG) translocon and subsequent membrane insertion ((37), Fig. 3A).

**Fig. 3.**
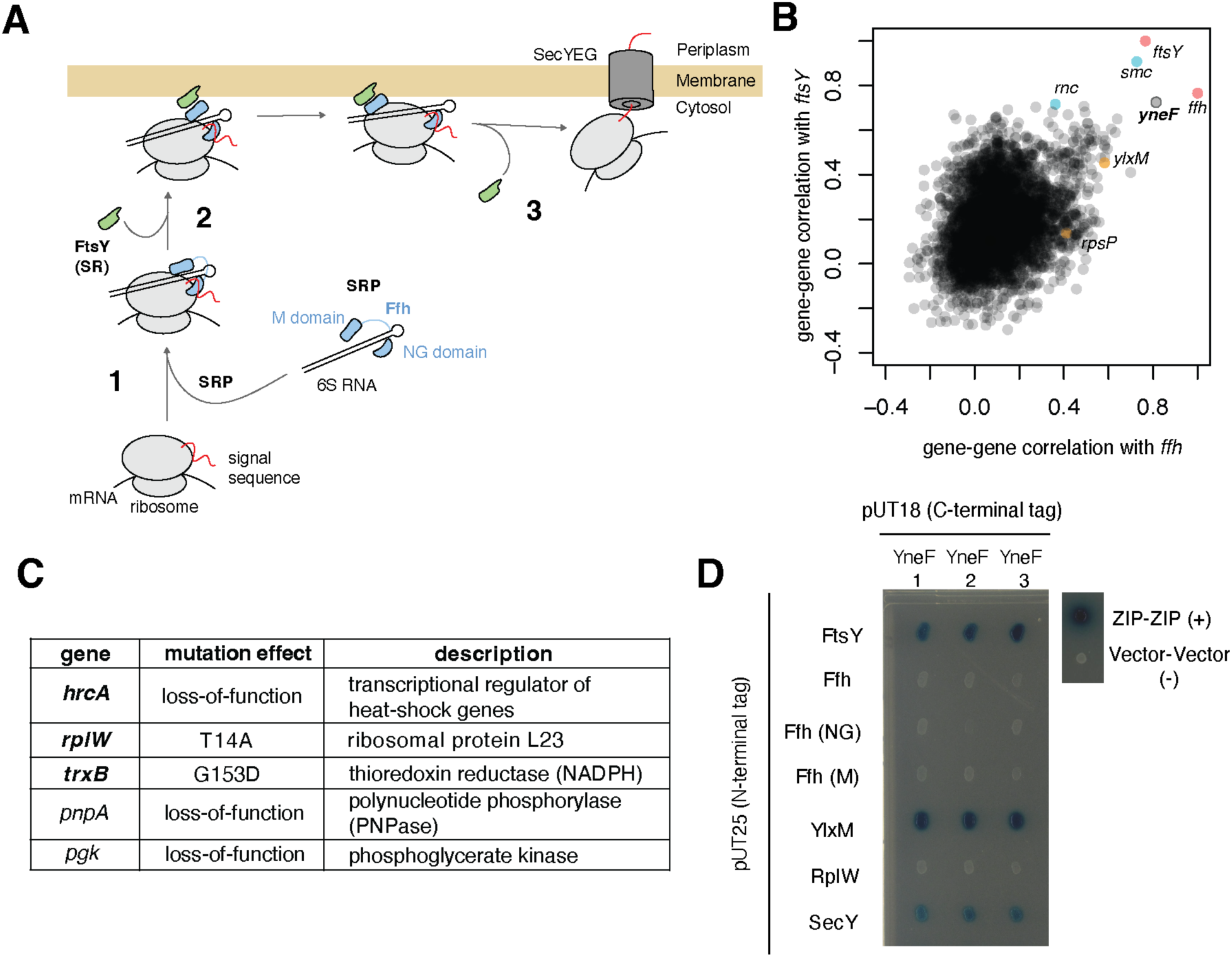
YneF is a novel, essential member of the SRP. A. SRP-mediated co-translational protein insertion. The SRP (Ffh and 6S RNA) binds to signal sequences as they are translated by the ribosome (1) and associates with FtsY (SR), which localizes the ribosome-SRP complex to the membrane (2), ultimately resulting in binding to the SecYEG translocon for protein insertion and disassembly of the complex (3). SecYEG also secretes non-SRP substrates. Schematic adapted from ref (37). B. Correlation values of each gene (n = 4162) with *ffh* are plotted against their correlation values with *ftsY* based on similarity of phenotypes across conditions. For genes with three gene knockdown bins, the maximum correlation is plotted. Cyan: genes sharing an operon with *ftsY* (*smc, rnc*). Orange: genes sharing an operon with *ffh* (*ylxM, rpsP, khpA*). Red: *ffh-ftsY* correlation. High correlation between *smc/rnc* and SRP genes is likely due to polar effects of their knockdown on *ftsY*. Highlighted genes with r ≈ 0 to *ffh* and *ftsY* may be occluded by other points. C. Suppressor mutations that rescued growth of a *yneF* knockout mutant on LB. Bolded genes are discussed in the text. Further information is provided in Table S4. D. Bacterial two-hybrid (BACTH) assay of YneF and other SRP components. Proteins are tagged with either the T18 or T25 domains of *B. pertussis* adenylate cyclase and three biological replicates are plated on minimal maltose media + Xgal (Methods, (46)). Positive (ZIP) and negative (vector) controls are shown. Blue color indicates a positive interaction.

Knockdown of *yneF* and other SRP components had strong positive S-scores in minimal media (MM), suggesting that this condition may rescue *yneF* deletion. To test this we transformed a previously designed *yneF* deletion construct (5) into WT *B. subtilis* and recovered transformants on MM but not on LB (Fig. S4B). This allowed us to identify suppressors of *ΔyneF* able to grow on LB. We streaked individual *ΔyneF* colonies growing on a MM plate onto an LB plate, selected colonies that grew, and performed whole genome sequencing. We identified 5 unique suppressors, three of which were consistent with YneF functioning in the SRP machinery (Fig. 3C, Table S3).

a. Deletion of *hrcA* suppressed *yneF* essentiality. HrcA represses two heat shock operons (*hrcA-grpE-dnaK-dnaJ-yqeT-yqeU-yqeV* and *groES-groEL*) and its deletion results in over-expression of these genes (38). Overexpression of *dnaK/J* is among the first cellular responses to SRP depletion (39) and suppresses translocon defects in *Escherichia coli* (40), likely by maintaining the folding state of substrates until they can enter the translocon. To test whether *ΔhrcA* suppresses *yneF* via overexpression of *dnaK/J,* we constructed double deletion mutants of *hrcA* and most genes in the *hrcA* operon (e.g., *ΔhrcA*/*ΔdnaK*) and tested whether these strains could be transformed with *ΔyneF*. We reasoned that if overexpression of an operon member was required for suppressing *yneF*, the triple deletion (e.g., *ΔhrcA*/*ΔdnaK/ΔyneF*) would not be viable. We were able to construct triple deletions with every gene in the *hrcA* operon except *ΔdnaJ* and *ΔdnaK*, indicating that deletion of *hrcA* rescues *ΔyneF* by overexpression of *dnaJ* and *dnaK* (Fig. S4C).
b. A point mutation (G153D) in *trxB,* which encodes thioredoxin reductase, suppressed *yneF* essentiality. We speculate that this mutation may enhance the TrxB thiol-dependent antioxidant system, protecting misfolded proteins from thiol oxidation.
c. A point mutation (T14A) in *rplW,* which encodes the essential L23 ribosomal protein, suppressed *yneF* essentiality. L23 plays a crucial role in the interaction between the ribosome and the SRP (41, 42) and the translocon (43–45), providing another link between YneF and the SRP pathway. Together, these suppressors support a role for YneF in the SRP pathway.

To better understand the function of YneF in the SRP machinery, we tested whether YneF associates with SRP components using an adenylate cyclase-based bacterial two-hybrid system in *E. coli* (BACTH; (46); Fig. 3D; Fig. S4D). We constructed N- and C- terminally tagged versions of YneF, FtsY, Ffh (the entire protein and NG- and M domains alone), RplW, SecY, and YlxM (a Firmicute-specific SRP accessory protein encoded upstream of and co-transcribed with *ffh* (47)), and tested the interactions of C- and N- terminally tagged YneF with all other proteins. N-terminally tagged YneF did not exhibit any interactions (Fig. S4D), likely because the N-terminal adenylate cyclase fragment prevents its proper membrane insertion. C-terminally tagged YneF interacted with FtsY (the SR), SecY, and YlxM (Fig. 3D). The interaction between YneF and the transmembrane components of the SRP (SecY and FtsY) is consistent with its predicted localization and establishes its involvement in the SRP. YneF also interacted with YlxM. YlxM modulates Ffh GTPase activity (47), raising the possibility that the interaction of these Firmicute-specific SRP components synchronize the cytoplasmic and membrane components of the SRP, perhaps working with the additional Alu domain of *B. subtilis* to effect developmental programs such as sporulation (48). Taken together, our phenotypic data, suppressor screen, and BACTH data firmly establish a role for YneF in the SPR machinery.

### YbfF drives resistance to β-lactams

The most successful and widespread class of antibiotics are those that target the cell wall, including β-lactams, which inhibit the transpeptidation (TP) of PG. To identify genes underpinning resistance to β-lactams, we searched for gene knockdowns with phenotypes specific to this class of drugs. Knockdowns of *ybfE* and *ybfF*, in the putative *ybfGFE* operon, were consistently, specifically, and exquisitely sensitized to β-lactams, but not other PG-targeting drugs such as the transglycosylase (TG) inhibitor moenomycin or the undecaprenyl pyrophosphate inhibitor bacitracin (Fig. 4A). Testing the phenotypes of single gene deletions (5) to deconvolute operon-level effects of CRISPRi knockdowns revealed that only *ΔybfF* was sensitized to β-lactams (Fig. S5A). *ybfF* encodes a unique polytopic membrane protein restricted to *B. subtilis* and a few closely related species (Fig. S5B). The *ΔybfF* mutant lysed in the presence of β-lactams, and this phenotype was rescued by complementation with *ybfF* (Fig. 4B). Importantly, overexpression of *ybfF* in a WT background resulted in increased resistance to β-lactams compared to a WT control (Fig. 4B), indicating that YbfF is a bona-fide β-lactam resistance mechanism.

**Fig. 4.**
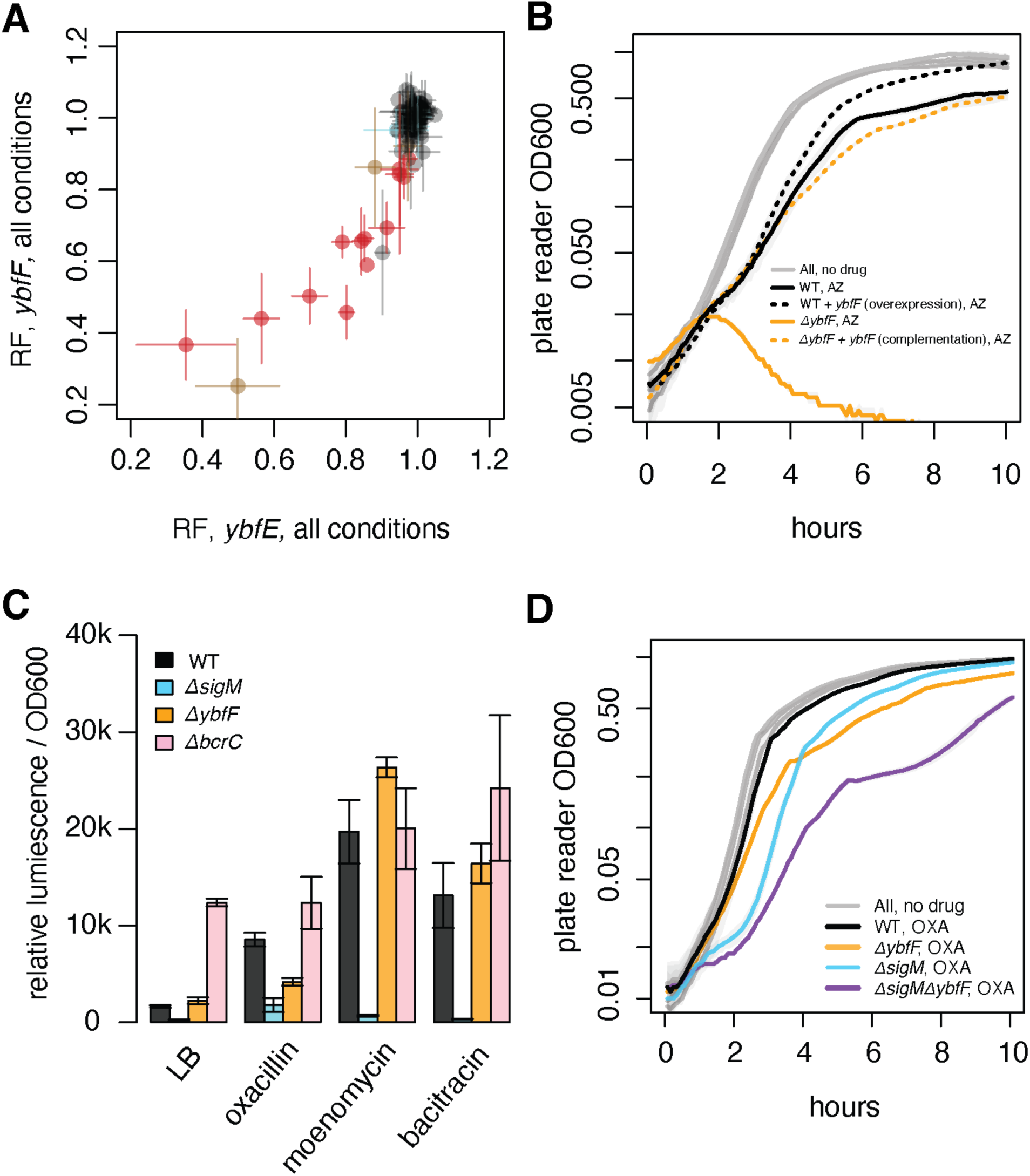
YbfF protects against β-lactam sensitivity in *B. subtilis*. A. Average RF values of *ybfE* and *ybfF* knockdown strains over all conditions (all concentrations of each stressor). Red = β-lactams; brown = EGCG, a catechin suggested to directly bind PG and cell surface proteins (88, 89); cyan = other cell wall antibiotics (moenomycin, bacitracin, cycloserine D, and fosfomycin; all with RFs ≈ 1.0) B. YbfF protects against β-lactams in a dose-dependent manner. WT (WT + P_xyl_-ybfF at *lac* without inducer), *ybfF* overexpression (WT + P_xyl_-ybfF at *lac* + 1% xylose induction), *ΔybfF* (*ΔybfF* + P_xyl_-ybfF without inducer), and *ybfF* complementation (*ΔybfF* + P_xyl_-ybfF + 1% xylose induction) were grown from a 1:100 dilution in LB only (grey, all strains) or LB + 6 μg/ml aztreonam (AZ). C. Maximum relative luminescence per OD600 during 4 hours of growth of WT, *ΔbcrC*, *ΔsigM*, or *ΔybfF* strains with a P_lux_-SigM reporter (Methods). Strains were diluted into LB, LB + 0.15μg/ml oxacillin, LB + 0.40μg/ml moenomycin, or LB + 40μg/ml bacitracin, or of a 96-well plate and OD600 and luminescence were measured. *ΔybfF* displays a depressed SigM response in response to the β-lactam oxacillin, but not the non-β-lactam antibiotics moenomycin or bacitracin. Full growth and luminescence curves provided in Fig. S6. D. *ΔsigMΔybfF* double deletion is more sensitive to β-lactams than single deletions alone. WT, *ΔybfF*, *ΔsigM*, and *ΔsigMΔybfF* were grown from a 1:100 dilution in LB only (grey, all strains) or LB + 0.04 μg/ml oxacillin (OXA).

Given the sensitivity of *ΔybfF* to β-lactams we next tested whether *ΔybfF* had an altered PG structure using HPLC ((49), Methods). However, the PG composition of *ΔybfF* and WT were similar (Fig. S5C), indicating that YbfF does not affect the composition of the PG in the absence of stress.

In our screen, SigM, which upregulates a large and diverse regulon in response to a variety of cell envelope stresses, was the primary sigma factor required to resist β-lactams ((50), Table S2). Using a previously described SigM reporter (51), we find that SigM induction in *ΔybfF* is reduced in response to the β-lactam oxacillin (*ΔybfF* maximum signal ∼0.48x that of WT; Fig. 4C, Fig. S6), but not in response to moenomycin and bacitracin (*ΔybfF* maximum signal ∼1.3x that of WT in both conditions; Fig. 4C, Fig. S6). However, this SigM induction defect is not solely responsible for β-lactam sensitivity of *ΔybfF*: a *ΔsigMΔybfF* mutant was significantly more sensitive to oxacillin than either single mutant (Fig. 4D), indicating that at least part of YbfF resistance to β-lactams is mediated by a SigM-independent pathway. Taken together, these data establish YbfF as a major factor for the resistance of *B. subtilis* to β-lactams and opens the door to further characterization of its mechanism.

### YybS is an ECF-transporter specificity subunit broadly implicated in cell wall homeostasis

We identified six *ΔybfF* colonies that grew within the zone of inhibition of aztreonam and subjected them to whole genome sequencing to identify suppressors. We found loss-of-function mutations in *ecfA_2_*, *ecfT,* and *yybS* (Table S3). *ecfA_2_* and *ecfT* encode components of the energy coupling factor (ECF) transporter, which binds reversibly to various specificity (S) subunits to import specific nutrients, metals, and vitamins into the cell (52, 53). *yybS* encodes a polytopic DUF2232-family protein. To determine if YybS is part of the ECF transporter, we used FoldSeek (54) to find structurally homologous proteins. YybS was similar to other DUF2232-family proteins (median e-value: 1.48e-12) and to ECF S-subunits predicted to transport riboflavin, queuosine, and other molecules (median e-value: 1.32e-2, Fig. S7A, Table S5). To further test whether YybS is an ECF S-subunit, we used AlphaFold3 to co-fold YybS with the ECF transporter complex (EcfT-EcfA_2_-EcfA_1_). AlphaFold3 predicted a high-confidence complex (overall ipTM = 0.84) in which YybS interacted with EcfT (ipTM = 0.81), EcfA_1_ (ipTM = 0.78), and EcfA_2_ (ipTM = 0.76) (Fig. 5A). YybS was predicted to bind in the same location and orientation as an S-subunit (e.g. PanU; RMSD = 0.607Å). A role for YybS as an ECF S-subunit is bolstered by the observation that some Actinobacterial DUF2232 domains are fused to transporter nucleotide binding domains like those of EcfA_1_/A_2_ or to EcfT homologs (e.g. A0A2U1E8B4 from *Actinomycetospora cinnamomea*).

**Fig. 5.**
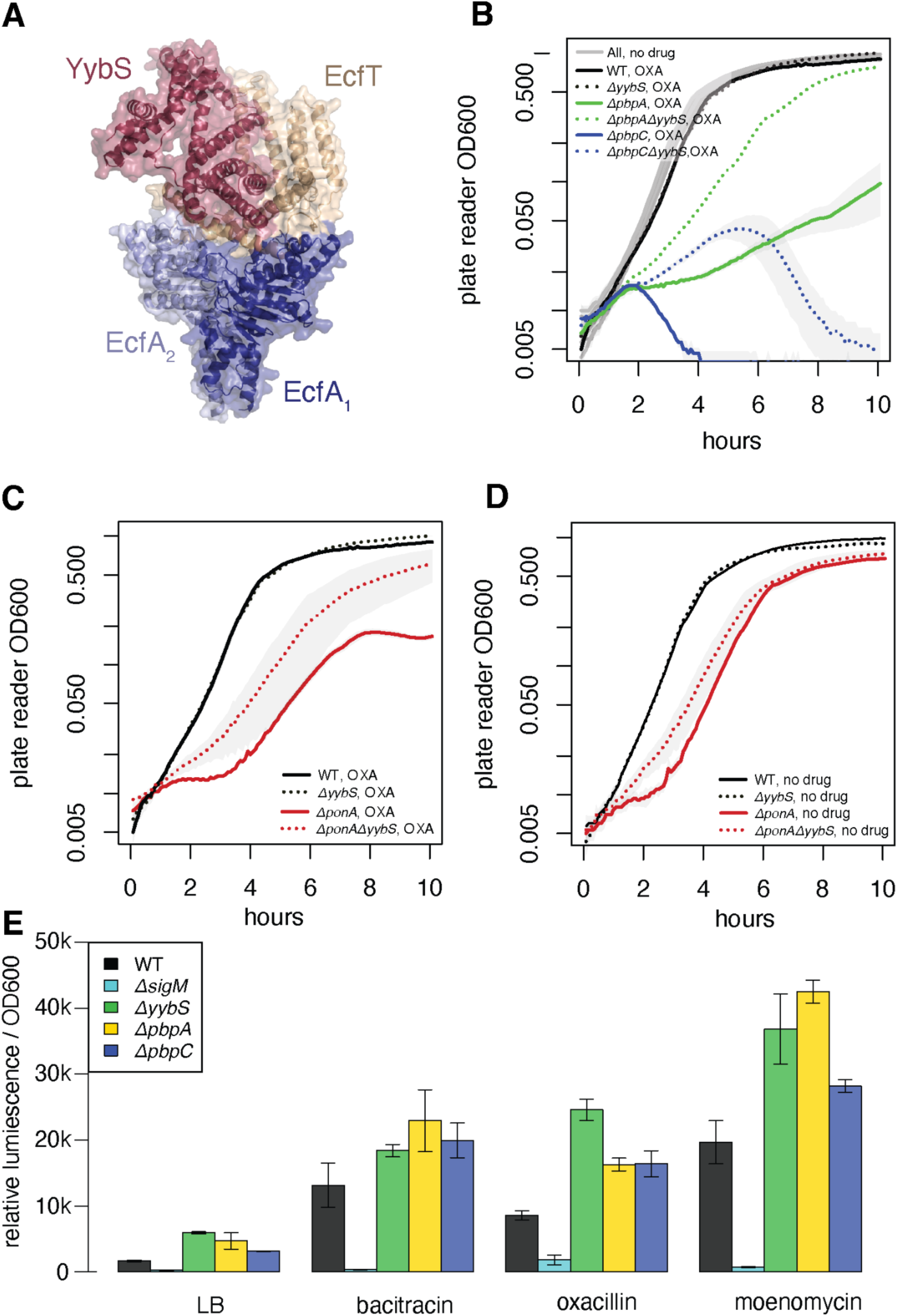
YybS is an ECF S-subunit which rescues β-lactam sensitivity in PBPs. A. Predicted AlphaFold3 structure of YybS, EcfA_1_, EcfA_2_, and EcfT (complex ipTM = 0.84) B. Deletion of *yybS* rescues β-lactam sensitivity of *ΔpbpC* and *ΔpbpA*. Overnight cultures were diluted 1:100 into LB or LB + 0.04 μg/ml oxacillin (OXA). C. Deletion of *yybS* rescues β-lactam sensitivity of *ΔponA*. Overnight cultures were diluted 1:100 into LB + 0.04 μg/ml oxacillin (OXA). D. Deletion of *yybS* enhances growth of *ΔponA*. Overnight cultures were diluted 1:100 into LB. E. Maximum relative luminescence per OD600 during 4 hours of growth of WT, *ΔsigM*, *ΔpbpA*, *ΔpbpC*, or *ΔyybS* strains with a P_lux_-SigM reporter (Methods). Strains were diluted into LB, LB + 40μg/ml bacitracin, or LB + 0.15μg/ml oxacillin of a 96-well plate and OD600 and luminescence were measured. *ΔyybS* displays a heightened SigM response compared to WT in all growth conditions, including LB. Full growth and luminescence curves provided in Fig. S6.

Although identified as *ΔybfF* suppressors, knockdown strains of *yybS*, *ecfA_1_, ecfA_2_,* and *ecfT* also exhibited subtle but significant resistance to cell wall targeting compounds in our screen (Fig. S7B-C), suggesting a role in cell wall homeostasis that is broader than *ybfF*. Indeed, in addition to rescuing the β-lactam sensitivity of *ΔybfF* (Fig. S7D), *ΔyybS* also rescued the β-lactam sensitivity of *ΔpbpA, ΔpbpC,* and *ΔponA* strains (Fig. 5B-C, Fig. S7D) and partially rescued the general growth defect of *ponA* (Fig. 5D). Because Δ*yybS* protected against the sensitivity of the PBP mutants, we tested whether *ΔyybS* had an altered PG structure using HPLC ((49), Methods). However, *ΔyybS* and WT exhibited similar PG structures (Fig. S5C), at least in the absence of stress.

We next tested whether *ΔyybS*-mediated protection might result, at least in part, from over-inducing SigM. Indeed, using the SigM reporter (51), we found that *ΔyybS* overinduces SigM in response to every tested cell wall drug (*ΔyybS* maximum signal ∼1.4-2.9x that of WT; Fig. 5E), and even slightly induces SigM in LB (*ΔyybS* maximum signal ∼3.6x that of WT; Fig. 5E), with maximal induction resembling the extent of induction in *ΔpbpA* or *ΔpbpC* strains (Fig. 5E; Fig. S6). Although the substrate imported by YybS has not yet been identified, these results suggest that in its absence, cell wall homeostasis is altered, leading to SigM overinduction. Together, these observations indicate that YybS plays a general role in maintaining cell wall homeostasis.

A role for YybS in the cell wall does not appear to be limited to *B. subtilis*; *S. aureus* transposon insertion mutants of *yybS* and *ecfT* also exhibited resistance to lysobactin, an antibiotic that binds to Lipid II (Fig. S7E-F). DUF2232-family proteins are widespread in firmicutes, actinobacteria, cyanobacteria, and alphaproteobacteria, and have been found to be essential in *Azospirillum brasilense* (an alphaproteobacteria) (55), *Synechococcus elongatus* (a cyanobacteria) (55), and *Clostridium difficile* (a firmicute) (56), suggesting that this previously unrecognized ECF S-subunit may influence cell wall integrity in diverse organisms response to various stressors.

### Adaptation to environmental change by community members driven by diverse response mechanisms

*B. subtilis* has evolved in the presence of numerous bacterial and fungal competitors (57). Conditioned media experiments, where one species is grown in the supernatant of another species, have been used to model interspecies interactions such as inhibition (58), cross-feeding (59, 60), or resource competition (61, 62). To understand the genetic underpinnings of *B. subtilis* competition with other species, we grew our library in the presence of conditioned media from a wide range of ecologically relevant organisms ((57), Methods). We identified three major sets of genes required for growth in the presence of spent media from multiple bacterial species.

First, knockdown of genes in the *mrp* operon, which encodes the essential Na+/H+ antiporter complex, strongly sensitized *B. subtilis* to conditioned media from *Klebsiella aerogenes, Shewanella oneidensis,* and *B. subtilis* strains (Fig. 6). Mrp antiporters (for “<u>M</u>ultiple resistance and <u>p</u>H”) play a role in resisting diverse stresses, particularly alkaline stress (63, 64), raising the possibility that those strains change the pH of the media. Consistent with this hypothesis, we found that the growth of *mrp* knockdown strains was strongly negatively correlated with the pH of the conditioned media used in the screen (r = -0.85 between pH and S-scores of medium-knockdown *mrpA* bin; Methods): *mrp* knockdown strains grew more poorly in more alkaline conditioned media. pH is a major driver of microbial distribution and success in the soil (65) and the increased sensitivity of *mrp* knockdown mutants in these conditioned media emphasizes the importance of maintaining pH homeostasis in the presence of other bacteria.

**Fig. 6.**
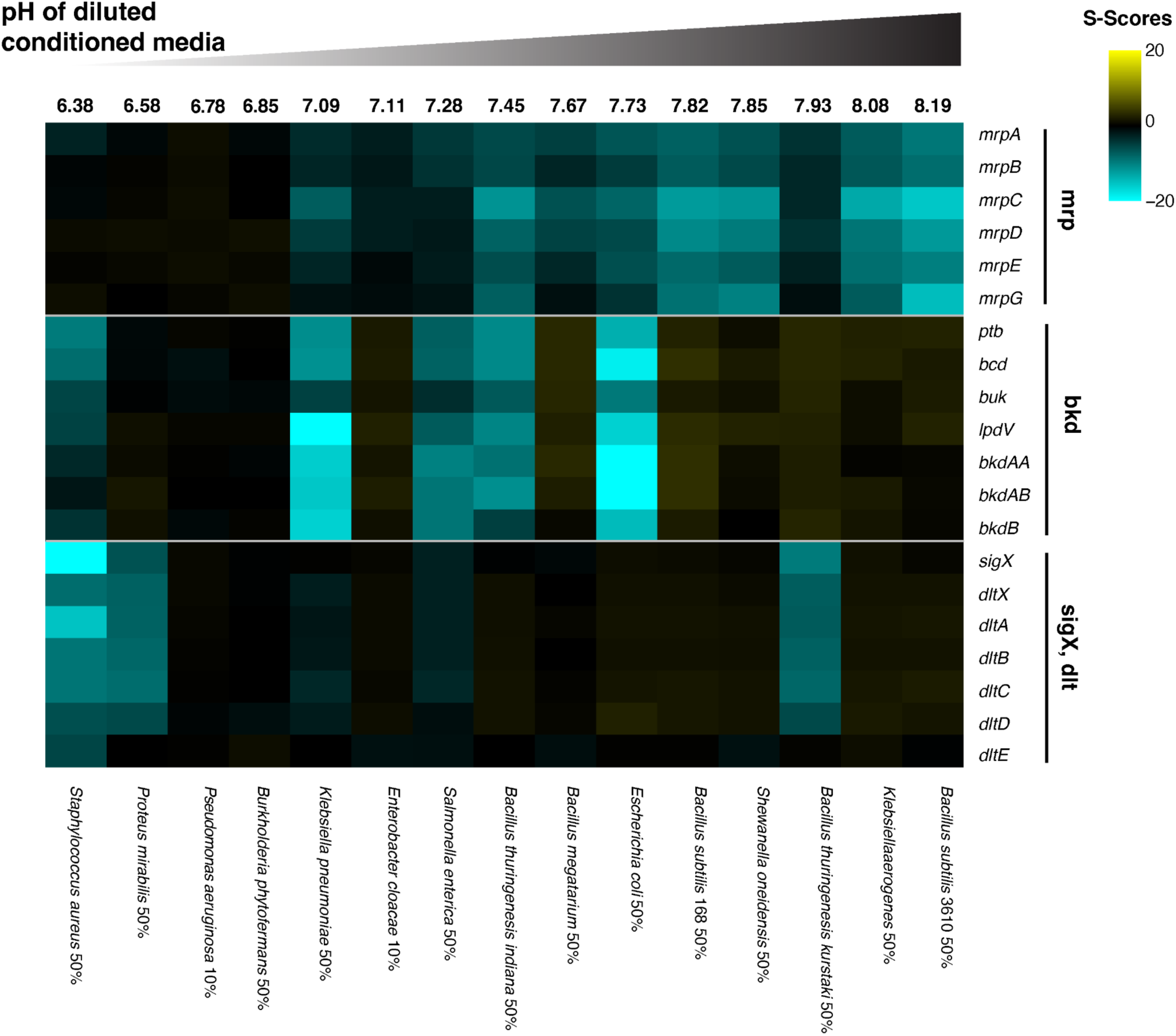
Major processes required for growth in diverse bacterial conditioned media. Heatmap of gene-chemical S-scores of operons required for growth in multiple bacterial conditioned media. Conditions (x-axis) are sorted by the pH of the conditioned media diluted with fresh LB (dilution percentage noted in labels, Methods). LB used in this screen is pH = 7.0. For genes targeted by multiple knockdown bins, only the most responsive bin (greatest sum of absolute S-scores across bacterial conditioned media) is shown.

Second, knockdown of the *bkd* operon (*ptb*-*bcd*-*buk*-*lpdV*-*bkdAA*-*bkdAB*-*bkdB,* hereafter *bkd*) sensitized cells to conditioned media from diverse species, including *E. coli, B. thuringiensis* var. Indiana, *Klebsiella pneumoniae,* and *S. aureus* (Fig. 6). *bkd* encodes the branched-chain keto acid dehydrogenase complex, which converts branched-chain amino acids to branched-chain fatty acids (BCFAs) (66), affecting membrane fluidity (67, 68). In *B. subtilis, bkd* is important for cold adaptation (69). We found that *bkd* is also important for growth in MM, glucose complete medium, and at pH 7.5 (Table S2), suggesting multiple ways in which *bkd* knockdowns could be sensitized to conditioned media. Our data implicate the production of BCFAs and, by extension, the alteration of membrane fluidity as an important aspect of interspecies competition.

Finally, knockdown strains of *sigX* (an ECF-type sigma factor) and the *dlt* operon (*dltXABCDE*, part of the SigX regulon (70)) were sensitized to conditioned media from *Proteus mirabilis, S. aureus*, and *B. thuringiensis* var. *Kurstaki* (Btk) (Fig. 6). SigX and Dlt are required for resistance to cationic antimicrobial peptides (CAMPs); SigX is activated by CAMPs and transcribes the *dlt* operon (70). Dlt lowers the net negative charge of the cell envelope by adding an alanine to the negatively-charged teichoic acids, rendering the cell less susceptible to CAMPs (70). CAMPs, such as bacteriocins, are widely produced by bacteria, including *S. aureus* and Btk (71–73). That *sigX* and *dlt* knockdown strains are sensitized to the conditioned media of specific species suggests that those species secrete relatively broad-spectrum CAMPs even when growing as a monoculture. Activation of the *dlt* operon may also be required to survive additional co-culture stressors beyond antimicrobials.

Together, these data demonstrate our screen captures responses to complex stress conditions and reveal major factors for resisting the environmental changes produced by the growth of other bacterial community members.

### Peptidoglycan stem peptides enable crossfeeding in gram-positive but not gram-negative bacteria

Cross-feeding enables the metabolic products of one bacterial species to sustain another one. These interactions are important for determining community composition and structure in the soil and the gut (60, 74, 75), but are incompletely characterized. We performed a global unbiased analysis of cross-feeding in *B. subtilis* by comparing the growth of our pooled knockdown library in MM (which enables crossfeeding) to the growth of an arrayed knockout library on plates of the same MM (5) (where colonies are too far apart to permit crossfeeding). We identified 5 knockdown strains (*gltA, gltB, gltC, icd,* and *alaT*) that are cross-fed (Fig. 7A).

**Fig. 7.**
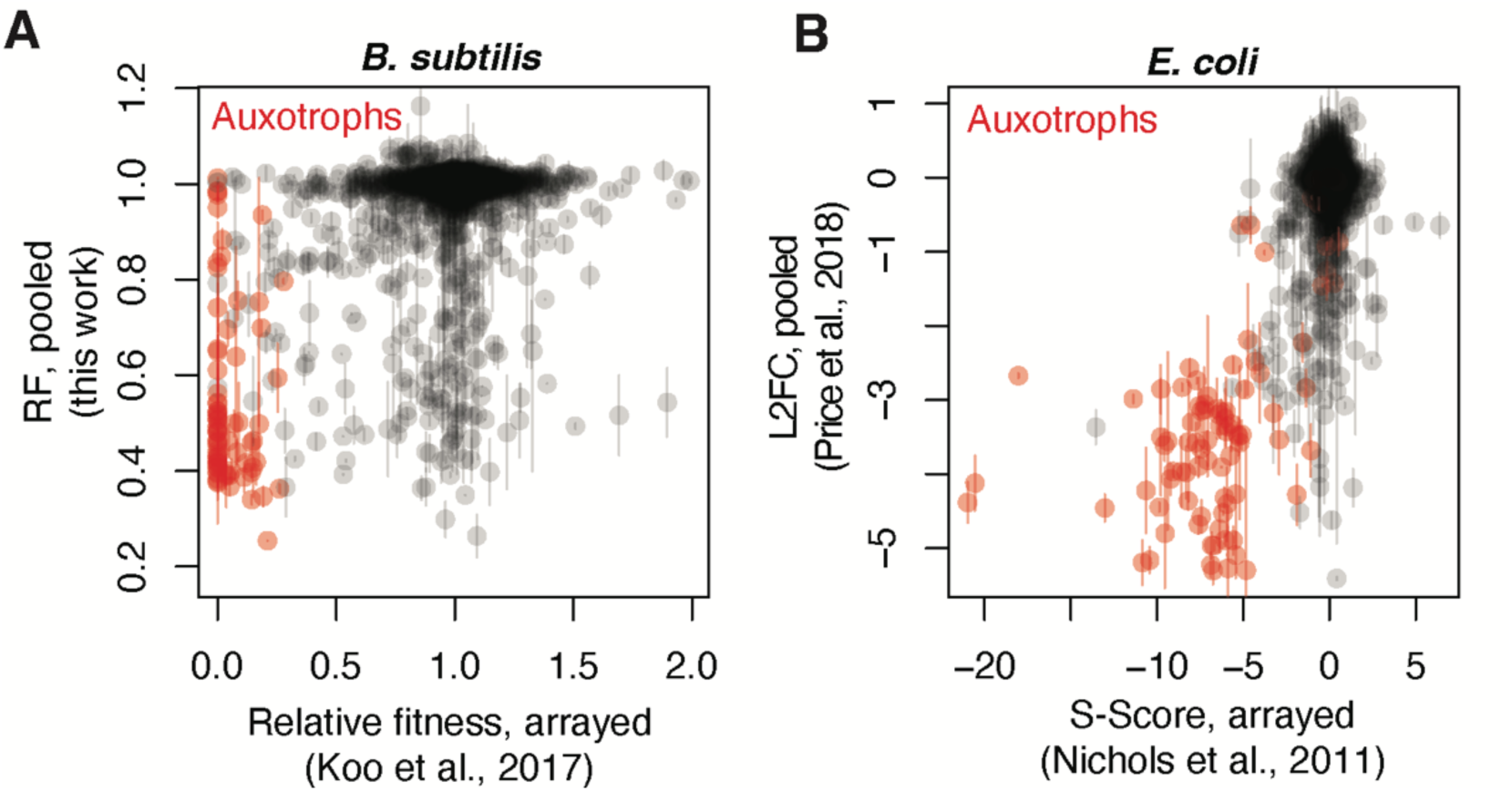
Cross-feeding in *B. subtilis* and *E. coli*. A. Growth of *B. subtilis* mutant strains grown either as single colonies of knockouts on MM agar plates (x-axis, (5)) or as a pooled knockdown library in MM (y-axis) (all genes present in both libraries; n = 3774 genes). For single colonies, WT growth is signified as a RF of 1.0. For growth of the library in liquid culture, WT growth is signified as a RF of 1.0. Error bars signify standard deviation of RF values in MM of the two targeting sgRNAs in pooled culture. Auxotrophs are highlighted in red (defined in ref (5)). Cross-feeding behavior of a strain would manifest as a RF of 0 as single colonies and a RF of 1.0 in pooled culture (upper left quadrant). B. Growth of *E. coli* knockout strains (90) grown as single colonies on MM agar plates (x-axis, (10)) or in pooled liquid MM culture growth (y-axis, (55)) (all genes present in both screens; n = 3410 genes). For single colonies, WT growth is signified as an S-score of 0. For growth in liquid culture, WT growth is signified as a L2FC of 0. Error bars signify standard deviation of two separate measurements of growth in pooled culture. Auxotrophs are highlighted in red (defined in ref (5)). Cross-feeding behavior of a strain would manifest as a negative S-score (i.e. S-score < -5) as single colonies and a L2FC of 0 in pooled culture (upper left quadrant).

To test whether this phenomenon represented true cross-feeding or was a byproduct of differences in growth between liquid and solid media (or between knockouts and knockdowns), we tested the ability of the *ΔgltA, ΔgltB,* and *ΔalaT* strains to grow on solid MM away from or in close proximity to WT *B. subtilis* using a previously developed cross-feeding assay (76). All three auxotrophic strains were able to grow on MM near (but not away from) WT. In contrast, non-cross-fed *ΔhisA* and *ΔcysJ* strains, which grew poorly on MM in both the arrayed- and pooled screen, were unable to grow on MM regardless of the presence of WT (Fig. S8), confirming the results of our analysis.

The *E. coli* genome-wide knockout library (77) has also been screened in MM in both pooled (55) and arrayed (10) settings. We therefore applied the same analysis to assess cross-feeding behaviour in *E. coli,* a gram-negative bacterium. Remarkably, we found no evidence of cross-feeding in *E. coli*: no mutants sick as single colonies showed growth akin to WT in pooled media (Fig. 7B), suggesting that *E. coli* is unable to support growth of any auxotrophs via cross-feeding. Because L-alanine, D-alanine, and D-glutamate are part of the *B. subtilis* and *E. coli* PG stem peptides (78), crossfeeding differences between gram-positive and gram-negative bacteria may be due to differences in PG recycling. Gram-positive bacteria perform multiple steps of PG recycling extracellularly, including the separation of the stem peptide (79–82), which may diffuse away and cross-feed auxotrophs or other organisms. In contrast, gram-negative bacteria retain over 90% of PG peptides within the periplasm for reuse (80, 83). More generally, gram-positive PG turnover and recycling may be an overlooked component of community-level nutrient metabolism.

## PERSPECTIVE

Here, we present a full-genome, CRISPRi-based chemical genomics screen in the model firmicute *B. subtilis*. By testing a large, diverse set of stressors (which increases the interpretability of correlated phenotypes), using CRISPRi instead of Tn-seq or knockout libraries (allowing us to query essential gene phenotypes), and optimizing the design of the library (for easy handling and cost-effective multiplexing), we make several important findings regarding *B. subtilis,* and, more broadly, gram-positive biology.

First, we identified roles for genes of unknown function, including essential genes. For example, we found that depletion of *yneF*, a highly expressed and deeply conserved gene with no annotated function, phenocopied depletion of SRP components *ffh* and *ftsY*. In follow-up experiments, we showed that YneF physically interacts with SRP components FtsY, SecY, and YlxM. Future studies will unravel the mechanistic role of YneF, and seek to understand why this auxiliary component became necessary in Firmicutes and Tenericutes. Our studies also identified new players in cell wall homeostasis. We showed that the uncharacterized Bacilli protein YbfF plays a major and specific role in resisting β-lactams. While investigating YbfF, we identified the DUF2232 protein YybS and ECF transporter components as suppressors of β-lactam sensitivity. Structural modeling revealed YybS to be a S-subunit for ECF transporters.

Despite the conserved structure of S-subunits, S-subunits importing distinct substrates have low sequence homology (53), making them difficult to annotate using sequence-based approaches. The specific substrate of YybS remains to be determined; future study of this S-subunit will clarify its role in cell wall homeostasis (84). Together, these results highlight the power of this approach to link genes to functions. Beyond these findings, our dataset contains many unexplored novel gene-chemical and gene-gene interactions, which will nucleate future discoveries.

Second, our dataset contains information on a large and diverse collection of antibiotics and antimicrobial compounds that cover a broad spectrum of targets and structural classes. This data can be leveraged to quickly determine the mechanism of action of novel antibiotics targeting gram-positive bacteria, as has been demonstrated for a transposon-based screen in *S. aureus* (11). Importantly, because our screen includes essential genes and substantially more chemicals, it can provide more accurate and targeted insights for a wider range of antimicrobials. Moreover, our data can be mined to provide insight into overlapping mechanisms of action of different compounds, potentially revealing synergistic antibiotic combinations (85).

Finally, we gained new insight into inter- and intra-species interactions. We tested our library in media conditioned by different species, allowing us to determine which genes are important for resisting environmental changes caused by bacterial competitors. Three processes were particularly important: pH homeostasis, modulation of membrane fluidity, and resistance to cationic antimicrobials. Within microbial communities, these factors are deeply intertwined (86, 87), and future studies combining a genomic approach with proteomics and/or proximity-based assays will provide deeper insight into these interactions. Our screen also uncovered a novel mechanism of intra-species crossfeeding. We found that *B. subtilis* (but not *E. coli*) is able to crossfeed glutamate and alanine, likely due to extracellular recycling of the stem peptide during PG turnover. The hypothesis that PG stem peptides are released by gram-positive (but not gram-negative) bacteria has major implications for our understanding of microbial community composition and structure in diverse niches.

In summary, our broad and comprehensive screen has revealed novel aspects of *B. subtilis* biology with implications for other gram-positive bacteria. These data will serve as a valuable resource for the *B. subtilis* research community, acting as a “hypothesis-generator” for future studies. Our approach, with its compact, information rich, sgRNA library design, acts as a model for future chemical genomics screens in other bacteria, which will bridge the gene-function gap and expand our knowledge of bacterial biology.

## ACKNOWLEDGEMENTS

We thank members of the Gross and Johnson labs for extensive helpful discussions. We thank Google DeepMind and Isomorphic Labs for providing free online access to AlphaFold3. SigM reporter strains were a gift from John Helmann. Bacterial two-hybrid strains were a gift from Michael Glickman. Yeast strains were a gift from Alexander Johnson. Work in the Gross lab was supported by the National Institutes of Health (NIH) grant R35 GM118061 (CAG) and by the Marcus Fund for Cell Health. Work in the Vollmer lab was supported by the University of Queensland. Sequencing was performed at the UCSF CAT, supported by UCSF PBBR, RRP IMIA, and NIH 1S10OD028511-01 grants.

## DATA AVAILABILITY

Raw sequencing data is deposited on NCBI-SRA Bioproject PRJNA1375834.

## TABLES WITH TITLES AND LEGENDS

**Table S1** RF for all sgRNAs across all conditions passing quality control.

**Table S2** S-scores for all genes across all conditions passing quality control.

**Table S3** Correlations between all genes across conditions.

**Table S4** Mapped suppressor mutations for *yneF* and *ybfF* knockout strains.

**Table S5** Foldseek e-values for YybS.

**Table S6** pH values of post-dilution bacterial conditioned media.

**Table S7** Information about primers, strains, and all stress conditions used.

**Data S1** Per-gene knockdown-fitness curves for all essential or impaired genes targeted by a mismatch sgRNA library in all conditions. Conditions with a combined |S| > 20 across sgRNA bins are highlighted.

## METHODS

### General strain manipulations and procedures

Unless otherwise noted, all strain construction and growth assays for *B. subtilis* were done in LB medium using antibiotics at the specified concentrations: Spectinomycin (100μg/ml), Chloramphenicol (6μg/ml), Kanamycin (7.5μg/ml). LB, glucose complete, and minimal media were prepared as previously described (Table S7, (5)).

### sgRNA plasmid construction

pJSHa77 (16) was modified to include a 6bp unique molecular identifier (UMI). pJSHa77 was digested with BamHI and an NNNNNN sequence was inserted into the BamHI digestion site with NEB HiFi (NEB #C2987) to create pJSHa77b.

### CRISPRi library construction

A *B. subtilis* CRISPRi library was designed and cloned as previously described (16), with minor modifications. Oligos encoding sgRNAs included ∼100bp homology with the pJSHa77b backbone were ordered from Twist Biosciences and amplified for 14 cycles. The pJSHa77b backbone was digested with BsaI-HFv2. The oligo library was inserted into the backbone via NEB HiFi and transformed into NEB 10beta *E. coli,* recovered, then grown overnight in 100mL LB and midiprepped the following day with the Qiagen Plasmid Plus kit (Qiagen #12945). The plasmid library was digested with NdeI for three hours at 37°C, cleaned up, and transformed into *B. subtilis* as previously described (16).

### sgRNA library design

The CRISPRi library targets the full genome: each essential gene was targeted by 12 sgRNAs with a range of predicted knockdown efficacies, and each non-essential gene was targeted by 2 fully complementary sgRNAs for a total library size of 11,559 sgRNAs. Selection of specific sgRNA categories are expanded below.

#### Essential Genes

The essential gene set was defined as the set of 300 genes previously investigated in ref (16). Using experimental RF data from ref (16), two of the 10 parent sgRNA families from each gene were selected from the original library. Parent sgRNA families were selected based on distance from the median experimental RF line. 5 of the 10 daughter mismatch sgRNAs of varying knockdown efficiencies were selected from each family based on bins and distance from the median as described in ref (16).

#### Nonessential Genes with Koo 2017 Phenotypes

17 nonessential gene knockouts in ref (5) had significant growth phenotypes in LB at 37°C. Of these, 10 were already included in ref (16) and the library was selected as described in “Essential Genes”. The remaining 7 (*mbl, lipL, qoxABCD,* and *rpmH*) were targeted by the two parent sgRNAs with the highest score and 10 mismatch daughters as described in ref (16, 17).

#### Nonessential Genes

For nonessential genes without strong phenotypes in ref (5) and passing exclusion criteria, sgRNAs were generated as described in ref (17) and scored based on observed knockdown efficacy weights (sequence logo in ref (17)). sgRNAs with more than 10 off-target hits and sgRNAs containing BsaI or NdeI restriction sites (used in cloning steps) were excluded. For each gene, the two highest-scoring sgRNAs were selected.

### Chemical genomics experiments and sequencing

Library aliquots were thawed, diluted to OD600 0.01, and grown in appropriate media (LB for all chemical stressors; MM, GC, or TSB if an alternative media was being tested) at 37°C to an OD600 of 0.10. From here, a t_0_ sample was taken, and the culture was back diluted to OD600 0.01 in 37°C growth media with the stressor and 1% xylose to induce dCas9 expression. When possible, two concentrations of each stressor were chosen for a 10% or 25% reduction in growth rate relative to WT *B. subtilis* in LB. After approximately 5 doublings, the culture was back diluted 1:10 into fresh media + 1% xylose and chemical stressor; after another 5 doublings, the culture was pelleted and frozen. OD600 was measured at each back-dilution and before collection via Synergy H4 plate reader (BioTek) calibrated to OD600 spectrophotometer readings. Genomic DNA was extracted from samples with the DNeasy Blood and Tissue Kit (Qiagen #69581) and libraries were prepared, pooled, and sequenced using Illumina NovaSeq as previously described (16).

### Relative fitness analysis and correction

RF values were calculated from sequence counts as previously described (16), with minor adjustments. Briefly, for each strain *X* with at least 10 counts at t_0_, we calculate the RF *F(x)* as:

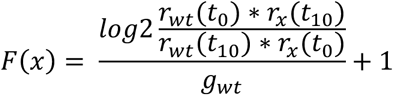

where *r_x_*(*t_0_)* is the fraction of strain *X* in the population at time *i* and *g_wt_* is the number of doublings of wildtype growth in the experiment. *g_wt_* is calculated from OD measurements. For this screen, *r_wt_(t_i_)* is calculated as the median value of nonessential sgRNAs.

Experimental samples with a median NE gene knockdown |L2FC| > 0.6 were discarded. Additionally, replicates with Pearson’s *r* < 0.8 with the other two replicates of the same condition were discarded. These criteria generally corresponded with bottleneck by low sample read depth.

Calculated RF values were then corrected for doubling artifacts as described previously (91). While we documented the number of doublings for each experiment, various technical errors and chemical effects can still preclude accurate doubling counts. Minimal and glucose complete media experiments were excluded from the correction, as there was no appropriate reference condition.

A small number of nonessential genes (<3%) were excluded from further analysis because the two sgRNAs targeting them had discrepant median RFs across experiments (at least one median RF > 0.9, difference in median RF > 0.1), indicating possible toxicity or off-target effects of one sgRNA and precluding a reliable phenotype. We took advantage of polar effects to salvage sgRNAs from a small number of genes: if one sgRNA had a high maximum correlation with another sgRNA targeting within the same operon (r > 0.5) and the other did not (maximum correlation with an operon-targeting sgRNA < 0.3), the highly correlated sgRNA was salvaged, and the other was discarded (Table S1). We note that several of the genes with discrepant sgRNAs share operons with essential genes, resulting in variable fitnesses due to polar effects.

### Calculation of chemical-gene S-scores

For nonessential genes, S-scores were calculated as described below. For essential genes and those targeted by a library of mismatch sgRNAs, sgRNAs were binned into three bins by strength (low, medium, and high knockdown efficiency), and S-scores for each bin were calculated separately.

RFs were transformed into S-scores as previously described (21). First, we took the median fitness of each gene knockdown across all conditions, and its dispersion about the median using the median absolute deviation (MAD) times 1.4826 to align with standard deviation. As applied previously, a floor was assigned to MADS; if a gene knockdown MAD was below the median MAD, it was replaced with the median MAD. Each RF was normalized by subtracting the gene knockdown RF median and scaled by the calculated MAD. For calculating S-scores, mean and variance for each gene-chemical interaction was calculated from all fitness values of relevant sgRNAs in each replicate of the given condition (e.g. for a nonessential gene targeted by two sgRNAs, in a condition with three technical replicates, mean and variance was calculated from the 6 RF measurements). These values were used for all gene-gene and condition-condition correlation analyses.

### Comparison to the STRING Database

Comparison to the STRING database was performed by downloading the *Bacillus subtilis* 168-specific interactions scores from http://string-db.org/ (92). Scores were culled to the “experimental” or “database” categories. Correlations between genes in the same operon were eliminated from the analysis.

### Disk Diffusion Assays

Disk diffusion assays were performed as described in (93). Briefly, 200uL of OD 1.0 culture was mixed with 4mL of 1% LB agar and evenly spread across an LB agar plate. 7mm filter discs were placed on the top agar and 3-5uL of antibiotic stock solution was applied to each disc.

### Suppressor Mapping

Suppressors of aztreonam sensitivity (Δ*ybfF*) or essentiality on LB (Δ*yneF*) were grown overnight in LB. Genomic DNA was extracted using the DNeasy Blood and Tissue Kit (Qiagen #69504). Whole-genome sequencing and variant calling was performed by SeqCenter in Pittsburgh, PA.

### AlphaFold3 Modelling

The webserver of AlphaFold3 was used to predict atomic models of the YneF dimer, YbfF dimer, and EcfA_1_/EcfA_2_/EcfT/YybS complex (94). Jobs were run with default options.

### Analysis of peptidoglycan composition

Cell wall purification and muropeptide analysis were performed as previously described in ref (49). Briefly, exponentially growing *B. subtilis* cells were harvested by centrifugation, the cell pellet resuspended in PBS and the suspension added dropwise into boiling 5% SDS solution. The crude cell walls were recovered by high-speed centrifugation, washed free of SDS, broken with glass beads in a Precellys homogenizer and incubated with DNase, RNase, and trypsin. The trypsin was inactivated by adding 1% SDS and incubating at 80°C. The cell walls were thoroughly washed with water and freeze-dried.

For removal of the wall teichoic acid, the cell wall was resuspended with 48% hydrofluoric acid, stirred at 4°C for 48 h and washed several times, resulting in purified peptidoglycan. The muropeptides (disaccharide peptide subunits) were released by digestion with the muramidase cellosyl (Hoechst, Germany), reduced with sodium borohydride (5 mg/ml in sodium borate buffer pH 9.0) and separated by HPLC (Shimadzu CBM 40) using a reversed phase Prontosil 120-3-C18-AQ 3 μM column (Bischoff) and a linear gradient from 100% solvent A (40 mM sodium phosphate, pH 4.5, 0.0003% sodium azide) to 100% solvent B (40 mM sodium phosphate, 20% methanol, pH 4.0) at 52°C, in 270 min. Muropeptides were detected at 202 nm and peaks identified by comparison with ref (49, 95).

### Bacterial Two-Hybrid (BACTH)

BACTH was performed as described in ref (46). Briefly, plasmids derived from pUT18 and pKT25 containing T18 and T25 fusions to *B. subtilis* proteins were transformed into a *cya-E. coli* strain and selected on LB agarose containing kanamycin (40µg/mL) and carbenicillin (100µg/mL). Resulting transformants were pinned on maltose minimal agarose containing kanamycin, carbenicillin, and X-gal (40µg/mL) and incubated for 2 days at 30°C. Growing on maltose minimal media provides a highly specific readout: only cells with a functional adenylate cyclase, resulting from association of T18 and T25 by the interaction of their two fused proteins, are able to utilize maltose and grow.

### SigM reporter assay

Luciferase reporter construction and measurement was performed as previously described (51). Luciferase measurements without antibiotic stress were performed by inoculating 1uL of mid-log cells into 99uL of fresh LB in a 96 well plate. Luciferase measurements with antibiotic stress were performed by adding 100uL of exponentially growing cells (OD ∼0.1) and the antibiotic to a 96 well plate. Plates were incubated at 37°C with shaking in a Spark Multimode Microplate reader (Tecan), and OD600 and luminescence were measured every 5 minutes. Promoter activity was normalized to “relative luminescence” by dividing the relative light units (RLU) by OD600. Assays were conducted with three biological replicates.

### High-throughput growth assays

All strains were grown to OD ∼1.0, then back-diluted 1:100 into 200 μL fresh LB (for induced growth curves, LB was supplemented with 1% xylose) and grown with shaking at 37°C in a Synergy H4 plate reader (BioTek) for 10 hours. OD600 was measured at 5-min intervals. Assays were conducted with three technical replicates unless otherwise noted.

### Conditioned media experiments

To make conditioned media, each species or strain was grown overnight in LB (bacteria) or TSB (yeast and a *B. subtilis* 168 control) at 37°C or 30°C to saturation. Cultures were spun down and supernatant was passed through a 0.22 μm PVDF filter to remove cells. Conditioned media was frozen at -80°C until use. For the screen, conditioned media was diluted to 50% or 10% final concentration with fresh media, whichever produced ∼90% WT growth rate of the library.

Growth assays of the library in conditioned media experiments were performed largely as described in “*Chemical genomics experiments and sequencing.”* Briefly, the library was thawed, diluted to OD600 0.01 and grown in appropriate media (LB or TSB) at 37°C to an OD600 of 0.10. A t_0_ sample was taken and the culture was back diluted to OD600 0.01 in 37°C diluted conditioned media with 1% xylose to induce dCas9 expression. After approximately 5 doublings, the culture was back diluted 1:10 into fresh diluted conditioned media + 1% xylose; after another 5 doublings, the culture was pelleted and frozen. Sample preparation was performed as described in “*Chemical genomics experiments and sequencing.”*

### Crossfeeding assays

Crossfeeding assays were performed as previously described (76). Briefly, a WT *B. subtilis* “donor” strain was struck onto a MM agar plate and incubated at 37°C. After 3 hours, an auxotrophic recipient strain was struck 3mm from the donor in parallel. Photos were taken after 48 hours of incubation.

---

**Fig. S1.**
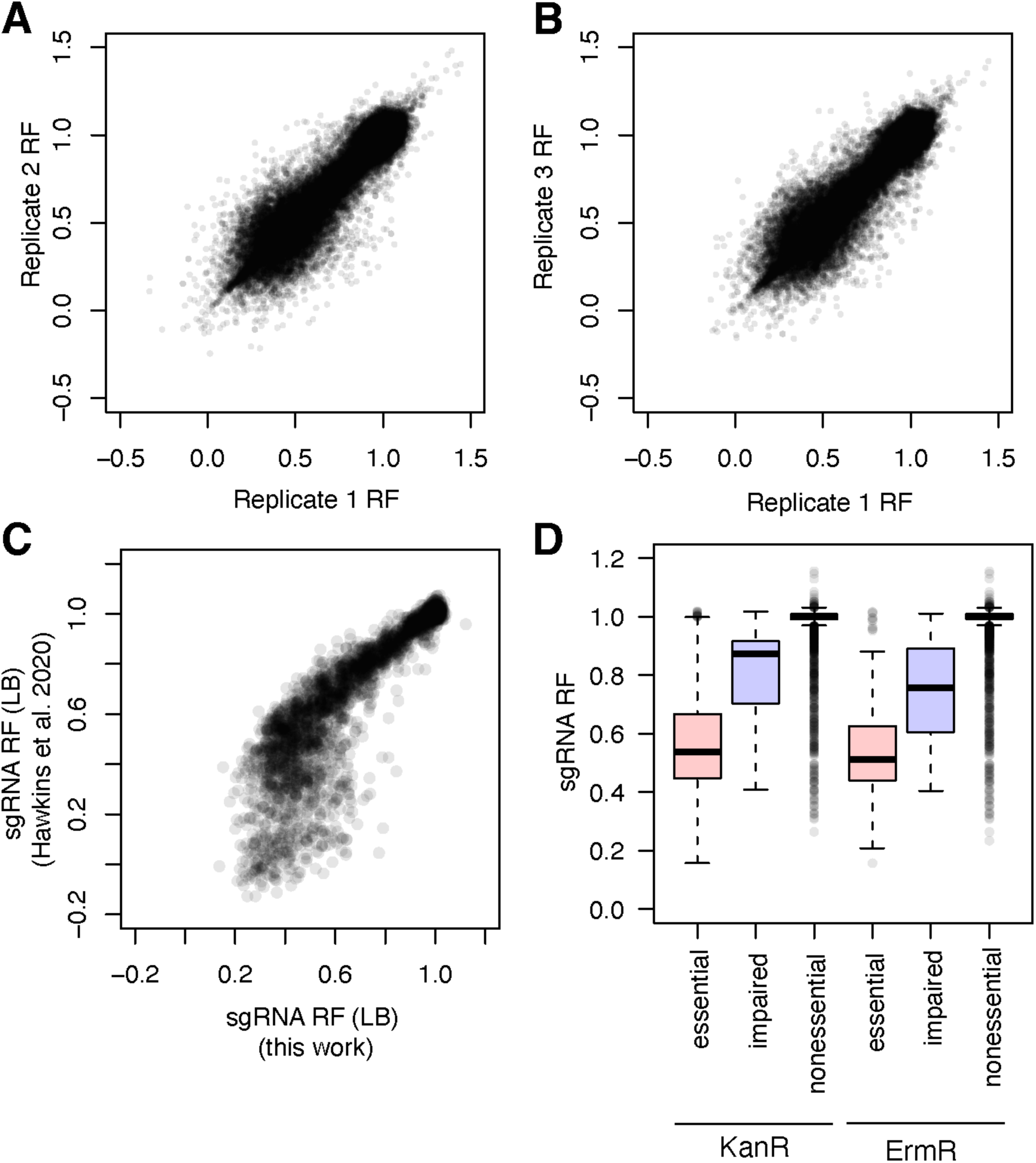
RF of sgRNAs is reproducible and consistent with previous measurements of sgRNA- or mutant fitness. A-B. RFs are reproducible across conditions. RF of all sgRNAs plotted pairwise from each experimental replicate of every condition (done in triplicate): 11,559 sgRNAs x 149 conditions ≈ 1.7 million RF values per replicate. Pearson’s *r* for all comparisons > 0.95. Condition replicates with severely abnormal median sgRNA behavior and/or outlier correlation values to the other triplicates generally indicate bottlenecking from read depth and were excluded during preprocessing prior to this analysis (Methods). 100,000 randomly chosen data points are displayed (out of approximately 1.7 million). C. RF of sgRNAs shared between our library and ref (16) are consistent. sgRNAs were selected from ref (16). Lower minimum RF in ref (16) reflects a higher sequencing depth in that study. Pearson’s *r* = 0.88. D. RF of fully-complementary sgRNAs targeting genes designated as “nonessential”, “impaired” or “essential” based on their single gene deletion phenotypes in KanR or ErmR libraries from ref (5). Box bounds: first and third quartile, midline: median, whiskers: most extreme datapoints. sgRNAs targeting nonessential genes with RF < 0.9 may reflect polar knockdown effects on essential operon partners.

**Fig. S2.**
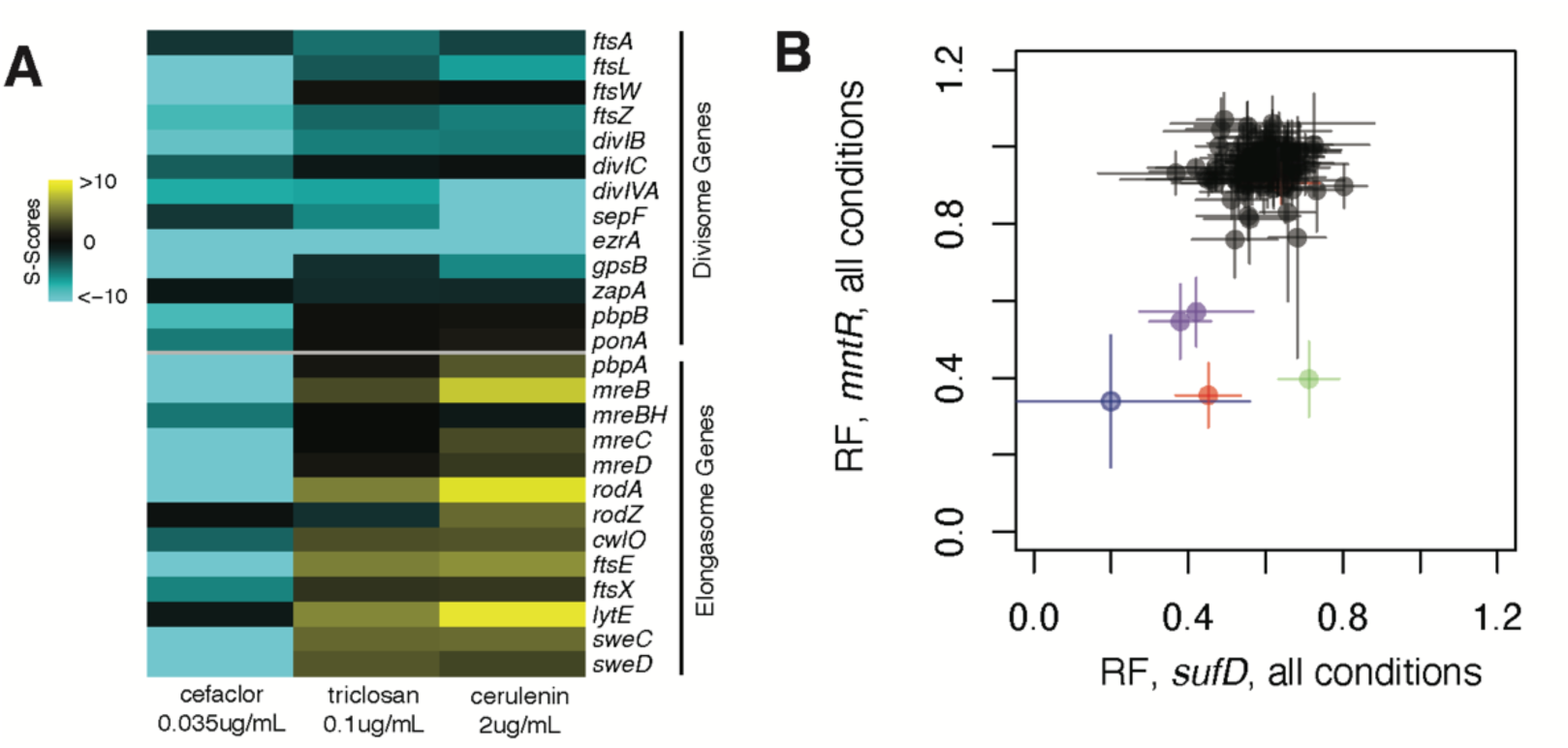
Expanded insight into previously identified pathway-pathway interactions. A. Gene-chemical S-Scores of genes involved in the elongasome or divisome against the cell wall drug cefaclor and the FAS inhibitors cerulenin and triclosan. As expected, knockdown of genes encoding the PG synthesis machineries of both the elongasome and divisome are sensitized to the β-lactam cefaclor. In contrast, inhibition of FAS by triclosan or cerulenin generally had a protective effect on knockdown of elongasome genes, but a sensitizing effect on knockdown of divisome genes. Drug concentrations with strongest S-scores are displayed. For genes targeted by multiple knockdown bins, only the most responsive bin (greatest sum of absolute S-scores across the drug concentrations used) is shown. B. RF values of *mntR* knockdown and *sufD* knockdown across all conditions. For *sufD*, the average RF of the two fully complementary sgRNAs is plotted. Red = mupirocin, Blue = dipyridyl (iron chelator), Green = excess manganese, Purple = minimal and glucose complete media.

**Fig. S3.**
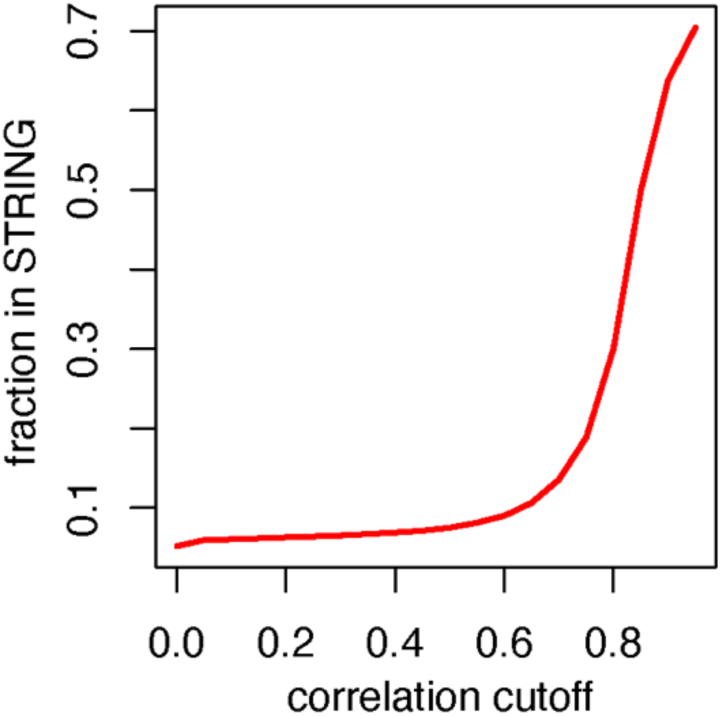
Correlations between the phenotypic profiles of genes reflect shared function. High-correlation gene pairs in the screen are enriched for STRING interactions. Intra-operon interactions (n=13k) are excluded due to CRISPRi polar effects. For genes with multiple knockdown bins, the maximum correlation between the bin sets was taken. STRING interaction scores are calculated from experimental and database datasets (Methods).

**Fig. S4.**
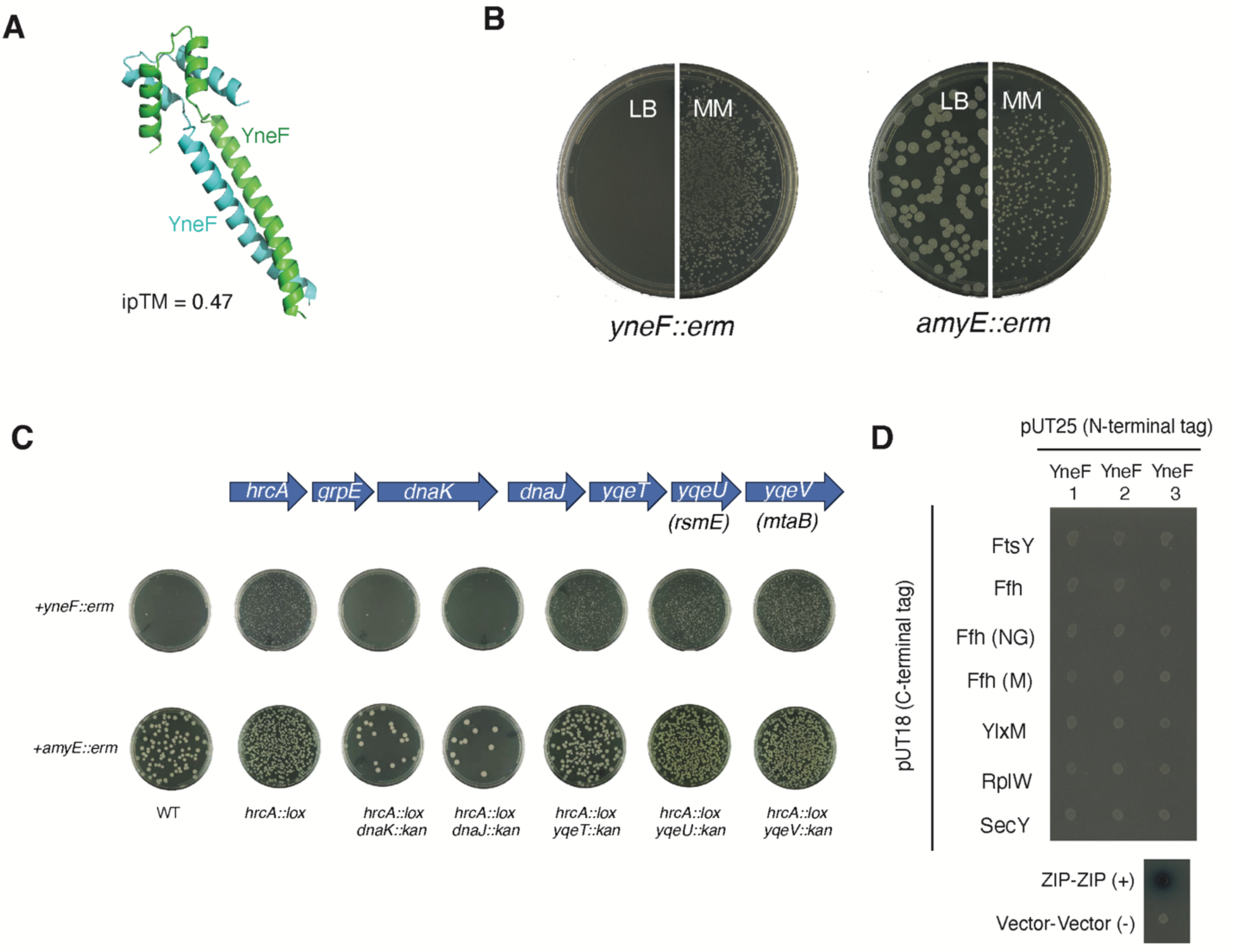
Additional data regarding YneF. A. Predicted YneF dimer by AlphaFold3 (ipTM = 0.47). B. Transformation of Δ*yneF::erm* or a Δ*amyE::erm* control into WT onto LB and MM. Plates were grown 72hr at 37°C before being photographed. Viable Δ*yneF::erm* transformants are produced when plating on MM but not LB. C. DnaKJ is required for suppression of *yneF* phenotype. Double deletions of *hrcA* and each gene in the *hrcA* operon were transformed with either Δ*yneF::erm* or an Δ*amyE::erm* control. Only the Δ*hrcA*Δ*dnaK* and Δ*hrcA*Δ*dnaJ* double deletions produced no viable colonies when transformed with Δ*yneF::erm*. We were not able to perform this assay with *groES-groEL* because they are essential. D. Bacterial two-hybrid (BACTH) assay of N-terminally tagged YneF and C-terminally tagged SRP components. Proteins are tagged with either the T18 or T25 domains of *B. pertussis* adenylate cyclase and plated on minimal maltose media + Xgal (Methods, (46)). Three biological replicates are shown. Improper insertion of YneF into the membrane due to the N-terminal tag likely precludes the positive interactions observed in C-terminally tagged YneF (Fig. 3D).

**Fig. S5.**
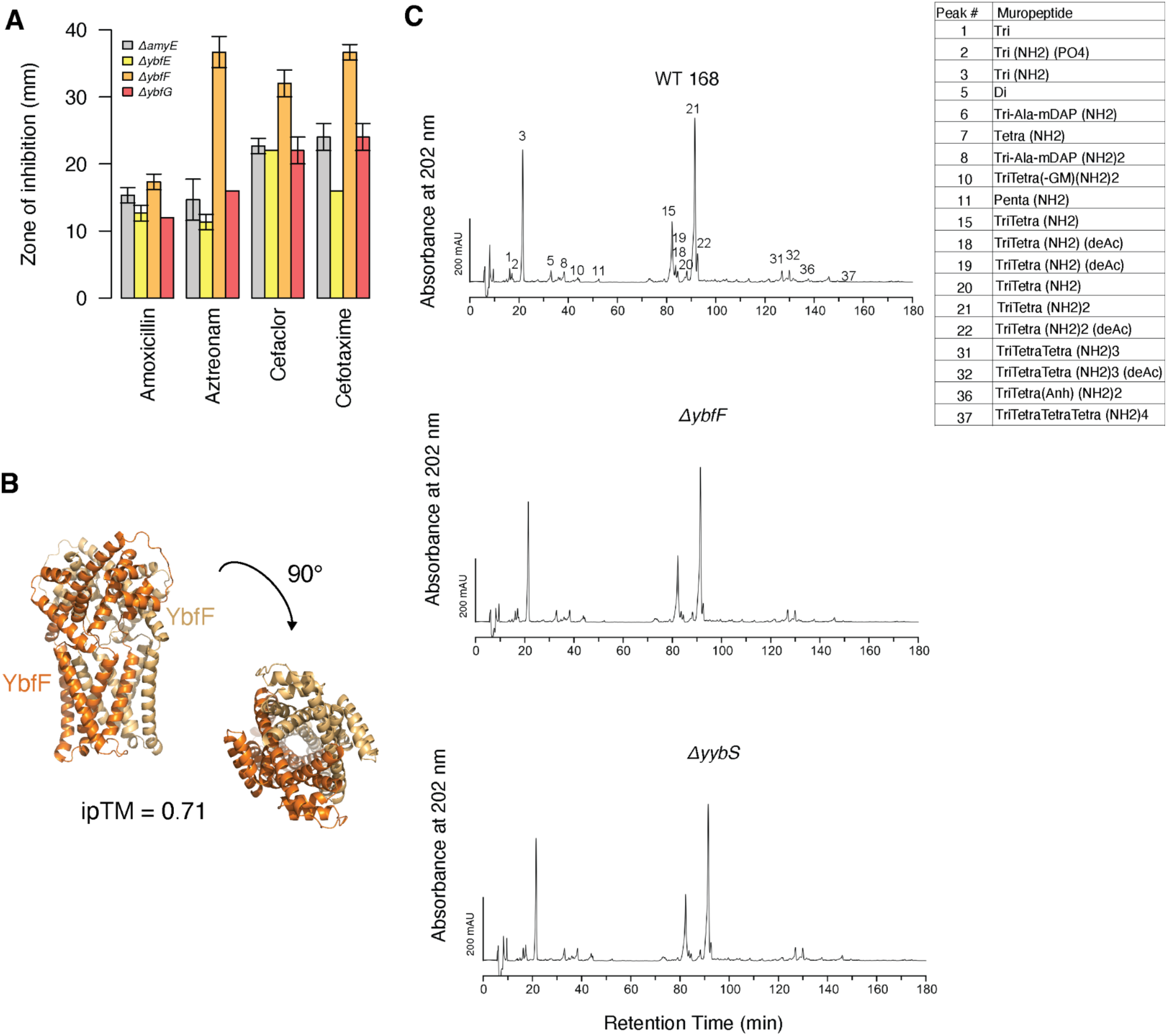
Additional data regarding *ybfF*, an uncharacterized gene with a role in cell wall homeostasis. A. Zone of inhibition from disk diffusion assays of *ΔybfE*, *ΔybfF, ΔybfG*, and an *ΔamyE* control against four β-lactams: 30μg aztreonam, 5μg cefaclor, 5μg amoxicillin, 5μg cefotaxime. Measurement is the diameter across the zone of inhibition (including filter disk). Results indicate *ybfF* is solely responsible for phenotypes. B. Predicted AlphaFold3 model of YbfF, a polytopic membrane protein. YbfF is predicted to act as a dimer (iptM = 0.71). C. Representative HPLC samples of muropeptides isolated from WT, *ΔybfF*, and *ΔyybS B. subtilis*.

**Fig. S6.**
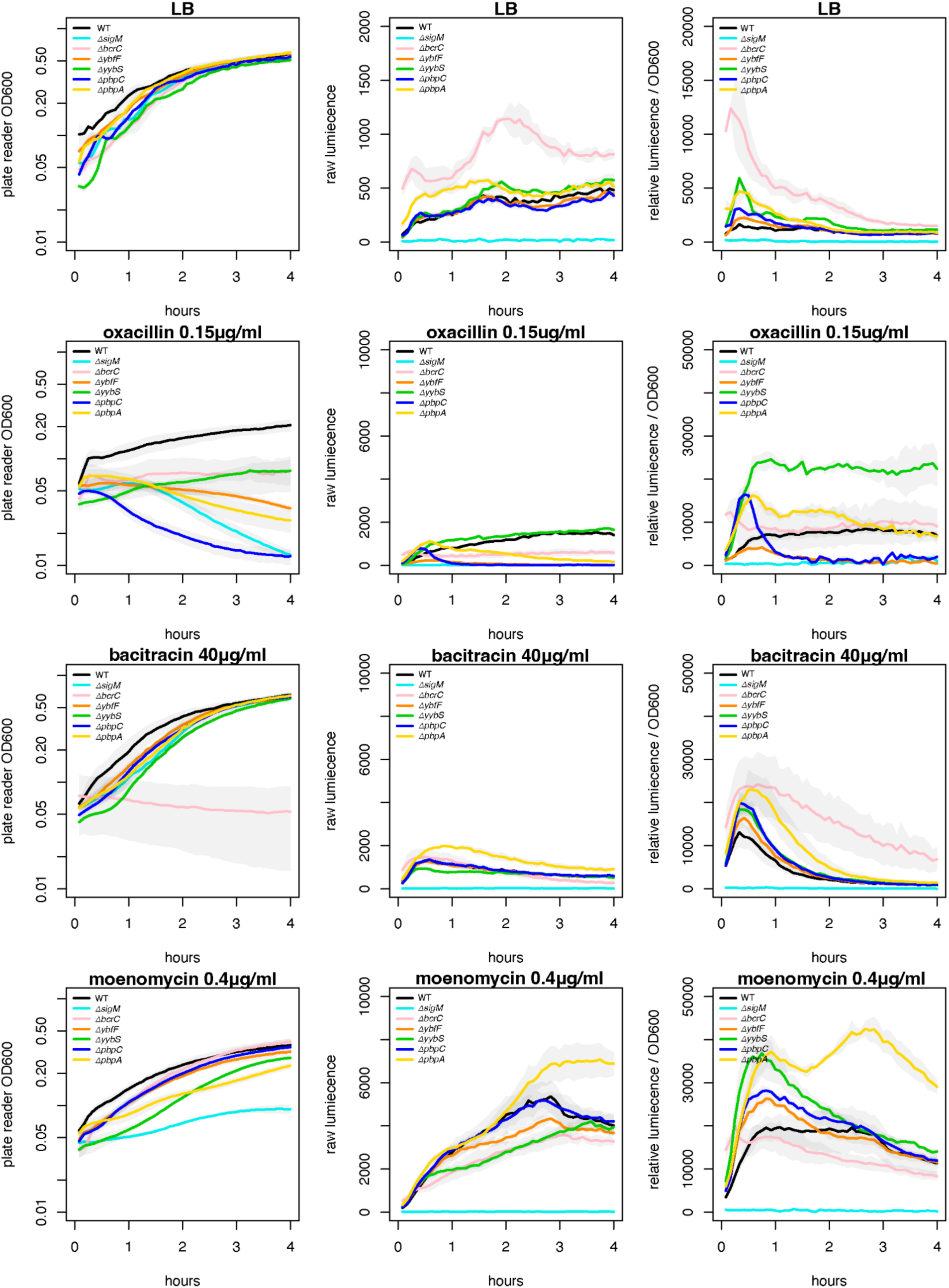
Expanded SigM reporter data. OD600, raw luminescence, and relative luminescence (raw luminescence / OD600) over 4 hours of growth of WT, *ΔbcrC*, *ΔsigM*, *ΔpbpA, ΔpbpC,* or *ΔybfF* mutants plus a P_lux_-SigM reporter (Methods). Strains were diluted into LB + 0.15μg/mL oxacillin, LB, LB + 40μg/mL bacitracin, or LB + 0.4μg/mL moenomycin in a 96-well plate and OD600 and luminescence were measured (Methods).

**Fig. S7.**
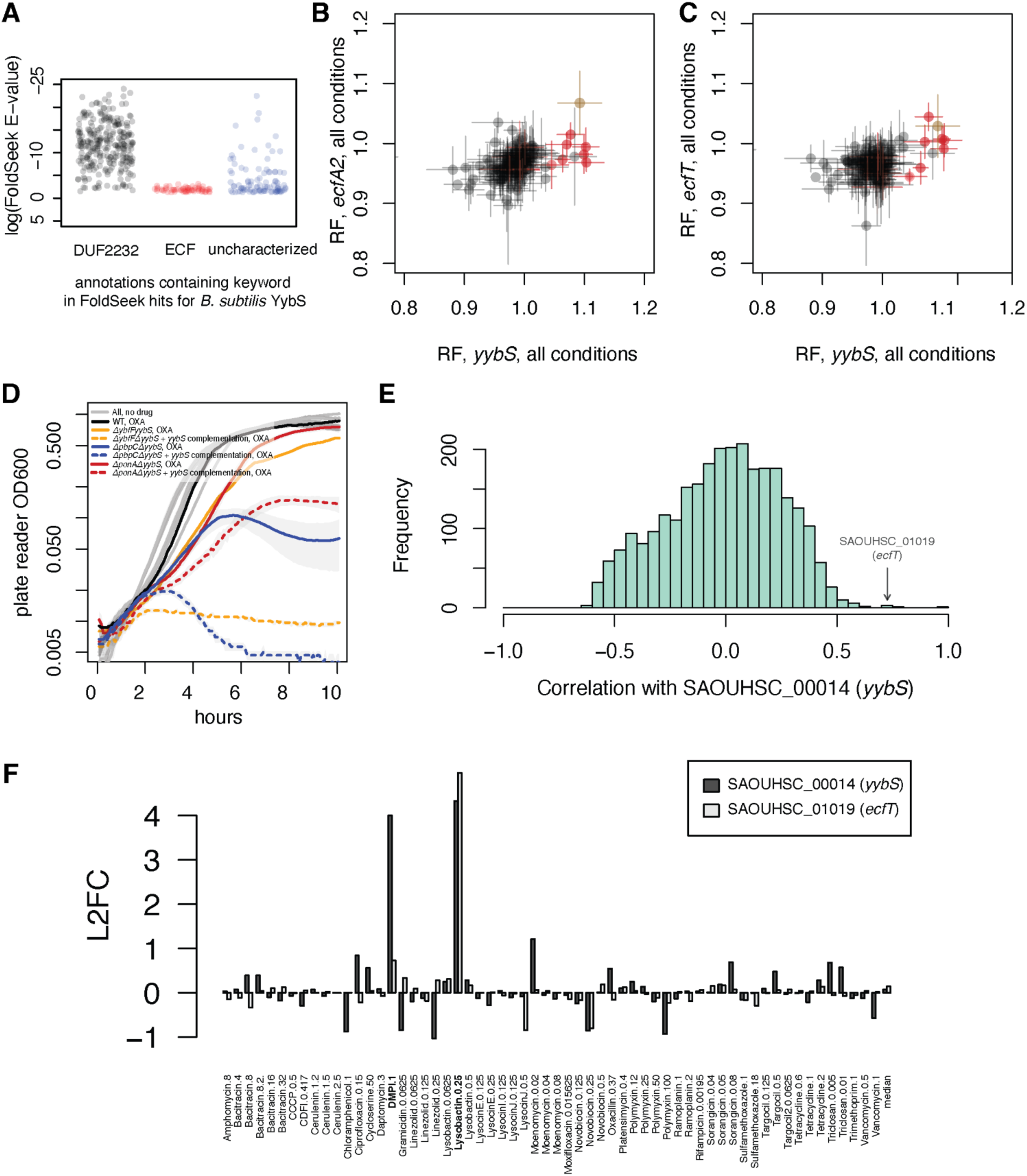
Additional data regarding YybS, a novel S-subunit of ECF transporters. A. FoldSeek results for the DUF2232 protein *B. subtilis* YybS grouped by annotation. FoldSeek was run with default parameters. B-C. Average RF values of *ecfA_2_, ecfT* and *yybS* knockdown strains over all conditions. Red = β-lactams; brown = ECGC. *yybS* shares an operon with the *gdpP*, which encodes a phosphodiesterase, and *rplI,* which encodes a nonessential large ribosomal protein; phenotypes may be due to polar effects. Novobiocin 3μg/ml was excluded from the plot due to a low *yybS* RF, likely due to polar effects. D. Deletion of *yybS* rescues β-lactam sensitivity of *ΔybfF* deletion and PBP deletions. This effect is reversed when *yybS* is exogenously expressed, indicating that this is due to the presence of YybS and not to a polar effect. Mid-log cultures of WT, *ΔybfFΔyybS* (*ΔybfFΔyybS* + P_xyl_-*yybS* without inducer), *ΔybfFΔyybS* + *yybS* complementation (*ΔybfFΔyybS* + P_xyl_-*yybS* + 1% xylose induction), *ΔpbpCΔyybS* (*ΔpbpCΔyybS* + P_xyl_-*yybS* without inducer)*, ΔpbpCΔyybS + yybS* complementation (*ΔpbpCΔyybS* + P_xyl_-*yybS* + 1% xylose induction), *ΔponAΔyybS* (*ΔponA* + P_xyl_-*yybS* without inducer), and *ΔponAΔyybS* (*ΔponA* + P_xyl_-*yybS* + 1% xylose induction) were diluted 1:100 in LB only (grey, all strains) or LB + 0.04 μg/ml oxacillin (OXA). E. Gene-gene correlations with *yybS* (SAOUHSC_00014) across all conditions in an *S. aureus* Tn-seq chemical genomics screen (11). *ecfT* (SAOUHSC_01019) is highly correlated with *yybS*, suggesting that *yybS* acts with ECF transporters in *S. aureus*. F. L2FC of *S. aureus* transposon deletions of *yybS* (SAOUHSC_00014) and *ecfT* (SAOUHSC_01019) across all conditions in a *S. aureus* Tn-seq chemical genomics screen (11). Deletion of *yybS* protects against lysobactin, which inhibits PG synthesis by binding Lipid II, and DPMI, which prevents export of Lipid II to the cell surface (11).

**Fig. S8.**
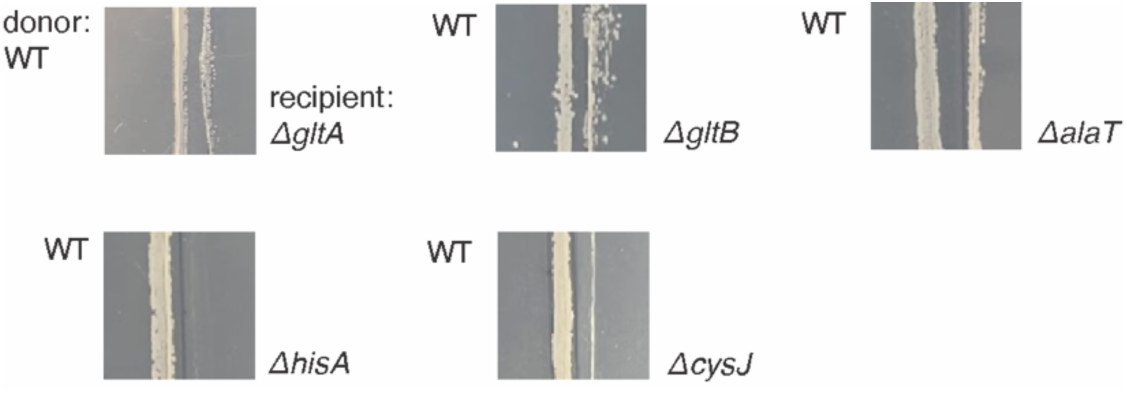
Crossfeeding of PG components. The “donor” strain (WT) was struck onto MM agar and incubated at 37°C. After 3 hours, the auxotrophic recipient strain was struck ∼3mm from the donor in parallel (Methods). Photos were taken after 48 hours of incubation. Growth of the auxotrophic recipient indicates crossfeeding.

## REFERENCES

1. K. Kobayashi, et al., Essential Bacillus subtilis genes. Proc. Natl. Acad. Sci. 100, 4678–4683 (2003).

2. P. Nicolas, et al., Condition-Dependent Transcriptome Reveals High-Level Regulatory Architecture in Bacillus subtilis. Science 335, 1103–1106 (2012).

3. C. M. Johnson, A. D. Grossman, Identification of host genes that affect acquisition of an integrative and conjugative element in Bacillus subtilis. Molecular Microbiology 93, 1284–1301 (2014).

4. J. M. Peters, et al., A Comprehensive, CRISPR-based Functional Analysis of Essential Genes in Bacteria. Cell 165, 1493–1506 (2016).

5. B.-M. Koo, et al., Construction and Analysis of Two Genome-Scale Deletion Libraries for Bacillus subtilis. Cell Syst 4, 291–305.e7 (2017).

6. F. J. O’Reilly, et al., Protein complexes in cells by AI-assisted structural proteomics. Mol. Syst. Biol. 19, e11544 (2023).

7. J. van Dijl, M. Hecker, Bacillus subtilis: from soil bacterium to super-secreting cell factory. Microbial Cell Factories 12, 3 (2013).

8. J. Stülke, A. Grüppen, M. Bramkamp, S. Pelzer, Bacillus subtilis, a Swiss Army Knife in Science and Biotechnology. J. Bacteriol. 205, e00102–23 (2023).

9. D. Wicke, J. Meißner, R. Warneke, C. Elfmann, J. Stülke, Understudied proteins and understudied functions in the model bacterium Bacillus subtilis—A major challenge in current research. Mol. Microbiol. 120, 8–19 (2023).

10. R. J. Nichols, et al., Phenotypic Landscape of a Bacterial *Cell*. Cell 144, 143–156 (2011).

11. M. Santiago, et al., Genome-wide mutant profiling predicts the mechanism of a Lipid II binding antibiotic. Nat Chem Biol 14, 601–608 (2018).

12. D. Leshchiner, et al., A genome-wide atlas of antibiotic susceptibility targets and pathways to tolerance. Nat. Commun. 13, 3165 (2022).

13. T. van Opijnen, K. L. Bodi, A. Camilli, Tn-seq: high-throughput parallel sequencing for fitness and genetic interaction studies in microorganisms. Nat. Methods 6, 767–772 (2009).

14. T. van Opijnen, A. Camilli, Transposon insertion sequencing: a new tool for systems-level analysis of microorganisms. Nat. Rev. Microbiol. 11, 435–442 (2013).

15. X. Liu, et al., High-throughput CRISPRi phenotyping identifies new essential genes in Streptococcus pneumoniae. Mol. Syst. Biol. 13, MSB167449 (2017).

16. J. S. Hawkins, et al., Mismatch-CRISPRi Reveals the Co-varying Expression-Fitness Relationships of Essential Genes in Escherichia coli and Bacillus subtilis. Cell Syst 11, 523–535.e9 (2020).

17. J. van Gestel, J. S. Hawkins, H. Todor, C. A. Gross, Computational pipeline for designing guide RNAs for mismatch-CRISPRi. Star Protoc 2, 100521 (2021).

18. M. R. Silvis, et al., Morphological and Transcriptional Responses to CRISPRi Knockdown of Essential Genes in Escherichia coli. mBio 12, 10.1128/mbio.02561-21 (2021).

19. B.-M. Koo, et al., Comprehensive double-mutant analysis of the Bacillus subtilis envelope using double-CRISPRi. bioRxiv 2024.08.14.608006 (2024). 10.1101/2024.08.14.608006.

20. N. Herrera, et al., The phenotypic landscape of the mycobacterial cell. bioRxiv 2025.11.14.688347 (2026). 10.1101/2025.11.14.688347.

21. S. R. Collins, M. Schuldiner, N. J. Krogan, J. S. Weissman, A strategy for extracting and analyzing large-scale quantitative epistatic interaction data. Genome Biol. 7, R63 (2006).

22. G. E. Schujman, K.-H. Choi, S. Altabe, C. O. Rock, D. de Mendoza, Response of Bacillus subtilis to Cerulenin and Acquisition of Resistance. J. Bacteriol. 183, 3032–3040 (2001).

23. R. Ohki, K. Tateno, T. Takizawa, T. Aiso, M. Murata, Transcriptional Termination Control of a Novel ABC Transporter Gene Involved in Antibiotic Resistance in Bacillus subtilis. J. Bacteriol. 187, 5946–5954 (2005).

24. J. M. Boyd, et al., Fpa (YlaN) is an iron(II) binding protein that functions to relieve Fur-mediated repression of gene expression in Staphylococcus aureus. mBio 15, e02310–24 (2024).

25. C. von Wachenfeldt, J. Hallgren, L. Hederstedt, YtkA (CtaK) and YozB (CtaM) function in the biogenesis of cytochrome c oxidase in Bacillus subtilis. Mol. Microbiol. 116, 184–199 (2021).

26. Y. R. Brunet, X. Wang, D. Z. Rudner, SweC and SweD are essential co-factors of the FtsEX-CwlO cell wall hydrolase complex in Bacillus subtilis. PLoS Genet. 15, e1008296 (2019).

27. R. H. Michna, F. M. Commichau, D. Tödter, C. P. Zschiedrich, J. Stülke, SubtiWiki–a database for the model organism Bacillus subtilis that links pathway, interaction and expression information. Nucleic Acids Res. 42, D692–D698 (2014).

28. B.-M. Koo, et al., Comprehensive genetic interaction analysis of the Bacillus subtilis envelope using double-CRISPRi. Cell Syst. 16, 101406 (2025).

29. F. Pompeo, et al., RagB stimulates the activity of the peptidoglycan polymerase RodA in Bacillus subtilis. EMBO Rep. 26, 4587–4606 (2025).

30. T. M. Bartlett, et al., FacZ is a GpsB-interacting protein that prevents aberrant division-site placement in Staphylococcus aureus. Nat. Microbiol. 9, 801–813 (2024).

31. J. R. Willdigg, Y. Patel, J. D. Helmann, A Decrease in Fatty Acid Synthesis Rescues Cells with Limited Peptidoglycan Synthesis Capacity. Mbio e00475–23 (2023). 10.1128/mbio.00475-23.

32. E. Michaud, et al., Stress-induced iron-sulfur cluster damage as a conserved trigger of the bacterial stringent response. Nat. Commun. (2026). 10.1038/s41467-026-70079-x.

33. X. Huang, J. Shin, A. Pinochet-Barros, T. T. Su, J. D. Helmann, Bacillus subtilis MntR coordinates the transcriptional regulation of manganese uptake and efflux systems. Mol. Microbiol. 103, 253–268 (2017).

34. J. I. Glass, et al., Essential genes of a minimal bacterium. Proc. Natl. Acad. Sci. 103, 425–430 (2006).

35. W. Liu, et al., Comparative Genomics of Mycoplasma: Analysis of Conserved Essential Genes and Diversity of the Pan-Genome. PLoS ONE 7, e35698 (2012).

36. M. Breuer, et al., Essential metabolism for a minimal cell. eLife 8, e36842 (2019).

37. M. M. Elvekrog, P. Walter, Dynamics of co-translational protein targeting. Curr. Opin. Chem. Biol. 29, 79–86 (2015).

38. A. Schulz, W. Schumann, hrcA, the first gene of the Bacillus subtilis dnaK operon encodes a negative regulator of class I heat shock genes. J. Bacteriol. 178, 1088–1093 (1996).

39. D. B. Bourgaize, et al., Loss of 4.5S RNA induces the heat shock response and lambda prophage in Escherichia coli. J. Bacteriol. 172, 1151–1154 (1990).

40. J. Wild, E. Altman, T. Yura, C. A. Gross, DnaK and DnaJ heat shock proteins participate in protein export in Escherichia coli. Genes & Development 6, 1165–1172 (1992).

41. S.-Q. Gu, F. Peske, H.-J. Wieden, M. V. Rodnina, W. Wintermeyer, The signal recognition particle binds to protein L23 at the peptide exit of the Escherichia coli ribosome. RNA 9, 566–573 (2003).

42. R. S. Ullers, et al., Interplay of signal recognition particle and trigger factor at L23 near the nascent chain exit site on the Escherichia coli ribosome. J. Cell Biol. 161, 679–684 (2003).

43. S. F. Ataide, et al., The Crystal Structure of the Signal Recognition Particle in Complex with Its Receptor. Science 331, 881–886 (2011).

44. F. Voigts-Hoffmann, et al., The Structural Basis of FtsY Recruitment and GTPase Activation by SRP RNA. Mol. Cell 52, 643–654 (2013).

45. K. Shen, et al., Molecular Mechanism of GTPase Activation at the Signal Recognition Particle (SRP) RNA Distal End*. J. Biol. Chem. 288, 36385–36397 (2013).

46. D. Ladant, Lipopolysaccharide Transport, Methods and Protocols. Methods Mol. Biol. 2548, 145–167 (2022).

47. M. L. Williams, P. J. Crowley, H. Adnan, B. L. Jeannine, YlxM Is a Newly Identified Accessory Protein That Influences the Function of Signal Recognition Particle Pathway Components in Streptococcus mutans. Journal of Bacteriology 196, 2043–2052 (2014).

48. M. Nishiguchi, K. Honda, R. Amikura, K. Nakamura, K. Yamane, Structural requirements of Bacillus subtilis small cytoplasmic RNA for cell growth, sporulation, and extracellular enzyme production. J. Bacteriol. 176, 157–165 (1994).

49. J. Sassine, J. Sousa, M. Lalk, R. A. Daniel, W. Vollmer, Cell morphology maintenance in Bacillus subtilis through balanced peptidoglycan synthesis and hydrolysis. Sci. Rep. 10, 17910 (2020).

50. W. Eiamphungporn, J. D. Helmann, The Bacillus subtilis σM regulon and its contribution to cell envelope stress responses. Mol. Microbiol. 67, 830–848 (2008).

51. H. Zhao, D. M. Roistacher, J. D. Helmann, Aspartate deficiency limits peptidoglycan synthesis and sensitizes cells to antibiotics targeting cell wall synthesis in Bacillus subtilis. Mol. Microbiol. 109, 826–844 (2018).

52. D. A. Rodionov, et al., A Novel Class of Modular Transporters for Vitamins in Prokaryotes. J. Bacteriol. 191, 42–51 (2008).

53. S. Rempel, W. K. Stanek, D. J. Slotboom, Energy-Coupling Factor–Type ATP-Binding Cassette Transporters. Annu. Rev. Biochem. 88, 1–26 (2018).

54. M. van Kempen, et al., Fast and accurate protein structure search with Foldseek. Nat. Biotechnol. 42, 243–246 (2024).

55. M. N. Price, et al., Mutant phenotypes for thousands of bacterial genes of unknown function. Nature 557, 503–509 (2018).

56. M. E. Alberts, et al., Analysis of essential genes in Clostridioides difficile by CRISPRi and Tn-seq. J. Bacteriol. 207, e00220–25 (2025).

57. H. Ma, D. Cornadó, J. M. Raaijmakers, The soil-plant-human gut microbiome axis into perspective. Nat. Commun. 16, 7748 (2025).

58. D. A. Schmitz, T. Wechsler, I. Mignot, R. Kümmerli, Predicting bacterial interaction outcomes from monoculture growth and supernatant assays. ISME Commun. 4, ycae045 (2024).

59. S. Rakoff-Nahoum, K. R. Foster, L. E. Comstock, The evolution of cooperation within the gut microbiota. Nature 533, 255–259 (2016).

60. G. D’Souza, et al., Ecology and evolution of metabolic cross-feeding interactions in bacteria. Nat. Prod. Rep. 35, 455–488 (2018).

61. Y. Wang, et al., Substrate Utilization and Competitive Interactions Among Soil Bacteria Vary With Life-History Strategies. Front. Microbiol. 13, 914472 (2022).

62. Y. Qiao, et al., Nutrient status changes bacterial interactions in a synthetic community. Appl. Environ. Microbiol. 90, e01566–23 (2023).

63. M. Ito, A. A. Guffanti, B. Oudega, T. A. Krulwich, mrp, a Multigene, Multifunctional Locus inBacillus subtilis with Roles in Resistance to Cholate and to Na+ and in pH Homeostasis. J. Bacteriol. 181, 2394–2402 (1999).

64. M. Ito, M. Morino, T. A. Krulwich, Mrp Antiporters Have Important Roles in Diverse Bacteria and Archaea. Front. Microbiol. 8, 2325 (2017).

65. H. K. Mod, et al., Predicting spatial patterns of soil bacteria under current and future environmental conditions. ISME J. 15, 2547–2560 (2021).

66. M. Debarbouille, R. Gardan, M. Arnaud, G. Rapoport, Role of BkdR, a Transcriptional Activator of the SigL-Dependent Isoleucine and Valine Degradation Pathway inBacillus subtilis. J. Bacteriol. 181, 2059–2066 (1999).

67. S. E. Diomandé, C. Nguyen-The, M.-H. Guinebretière, V. Broussolle, J. Brillard, Role of fatty acids in Bacillus environmental adaptation. Front. Microbiol. 6, 813 (2015).

68. M. Gohrbandt, et al., Low membrane fluidity triggers lipid phase separation and protein segregation in living bacteria. EMBO J. 41, EMBJ2021109800 (2022).

69. T. Kaan, G. Homuth, U. Mäder, J. Bandow, T. Schweder, Genome-wide transcriptional profiling of the Bacillus subtilis cold-shock response. Microbiology 148, 3441–3455 (2002).

70. M. Cao, J. D. Helmann, The Bacillus subtilis Extracytoplasmic-Function σX Factor Regulates Modification of the Cell Envelope and Resistance to Cationic Antimicrobial Peptides. J. Bacteriol. 186, 1136–1146 (2004).

71. E. L. Salazar-Marroquín, L. J. Galán-Wong, V. R. Moreno-Medina, M. Á. Reyes-López, B. Pereyra-Alférez, Bacteriocins synthesized by Bacillus thuringiensis. Rev. Méd. Microbiol. 27, 95–101 (2016).

72. L. L. Newstead, K. Varjonen, T. Nuttall, G. K. Paterson, Staphylococcal-Produced Bacteriocins and Antimicrobial Peptides: Their Potential as Alternative Treatments for Staphylococcus aureus Infections. Antibiotics 9, 40 (2020).

73. I. Sugrue, R. P. Ross, C. Hill, Bacteriocin diversity, function, discovery and application as antimicrobials. Nat. Rev. Microbiol. 22, 556–571 (2024).

74. S. Starke, et al., Amino acid auxotrophies in human gut bacteria are linked to higher microbiome diversity and long-term stability. ISME J. 17, 2370–2380 (2023).

75. E. J. Culp, A. L. Goodman, Cross-feeding in the gut microbiome: Ecology and mechanisms. Cell Host Microbe 31, 485–499 (2023).

76. K. R. Sidiq, M. W. Chow, Z. Zhao, R. A. Daniel, Alanine metabolism in Bacillus subtilis. Mol. Microbiol. 115, 739–757 (2021).

77. T. Baba, et al., Construction of Escherichia coli K-12 in-frame, single-gene knockout mutants: the Keio collection. Mol. Syst. Biol. 2, MSB4100050 (2006).

78. W. Vollmer, D. Blanot, M. A. D. Pedro, Peptidoglycan structure and architecture. FEMS Microbiol. Rev. 32, 149–167 (2008).

79. J. Mauck, L. Glaser, Turnover of the cell wall of bacillus subtilis W-23 during logarithmic growth. Biochem. Biophys. Res. Commun. 39, 699–706 (1970).

80. M. C. Gilmore, F. Cava, Bacterial peptidoglycan recycling. Trends Microbiol. (2024). 10.1016/j.tim.2024.11.004.

81. J. Mauck, L. Chan, L. Glaser, Turnover of the Cell Wall of Gram-positive Bacteria. J. Biol. Chem. 246, 1820–1827 (1971).

82. S. Litzinger, et al., Muropeptide Rescue in Bacillus subtilis Involves Sequential Hydrolysis by β-N-Acetylglucosaminidase and N-Acetylmuramyl-l-Alanine Amidase. J. Bacteriol. 192, 3132–3143 (2010).

83. E. W. Goodell, U. Schwarz, Release of cell wall peptides into culture medium by exponentially growing Escherichia coli. J. Bacteriol. 162, 391–397 (1985).

84. M. Partipilo, D. J. Slotboom, The S-component fold: a link between bacterial transporters and receptors. Commun. Biol. 7, 610 (2024).

85. E. Cacace, et al., Systematic analysis of drug combinations against Gram-positive bacteria. Nat. Microbiol. 8, 2196–2212 (2023).

86. G. Orazi, K. L. Ruoff, G. A. O’Toole, Pseudomonas aeruginosa Increases the Sensitivity of Biofilm-Grown Staphylococcus aureus to Membrane-Targeting Antiseptics and Antibiotics. mBio 10, 10.1128/mbio.01501-19 (2019).

87. A. Aranda-Díaz, et al., Bacterial interspecies interactions modulate pH-mediated antibiotic tolerance. eLife 9, e51493 (2020).

88. W.-H. Zhao, Z.-Q. Hu, S. Okubo, Y. Hara, T. Shimamura, Mechanism of Synergy between Epigallocatechin Gallate and β-Lactams against Methicillin-Resistant Staphylococcus aureus. Antimicrob. Agents Chemother. 45, 1737–1742 (2001).

89. M. Nakayama, et al., Mechanism for the antibacterial action of epigallocatechin gallate (EGCg) on Bacillus subtilis. *Biosci., Biotechnol.*, Biochem. 79, 845–854 (2015).

90. V. de Berardinis, et al., A complete collection of single-gene deletion mutants of Acinetobacter baylyi ADP1. Mol. Syst. Biol. 4, MSB200810 (2008).

91. L. M. Kim, H. Todor, C. A. Gross, Correction of a widespread bias in pooled chemical genomics screens improves their interpretability. Mol. Syst. Biol. 20, 1173–1186 (2024).

92. D. Szklarczyk, et al., STRING v10: protein–protein interaction networks, integrated over the tree of life. Nucleic Acids Res. 43, D447–D452 (2015).

93. Y. Luo, J. D. Helmann, Analysis of the role of Bacillus subtilis σM in β-lactam resistance reveals an essential role for c-di-AMP in peptidoglycan homeostasis. Mol. Microbiol. 83, 623–639 (2012).

94. J. Abramson, et al., Accurate structure prediction of biomolecular interactions with AlphaFold 3. Nature 630, 493–500 (2024).

95. A. Atrih, G. Bacher, G. Allmaier, M. P. Williamson, S. J. Foster, Analysis of Peptidoglycan Structure from Vegetative Cells of Bacillus subtilis 168 and Role of PBP 5 in Peptidoglycan Maturation. J. Bacteriol. 181, 3956–3966 (1999).

